# *Xist* Deletion in B Cells Results in Systemic Lupus Erythematosus Phenotypes

**DOI:** 10.1101/2024.05.15.594175

**Authors:** Claudia D. Lovell, Nikhil Jiwrajka, Hayley K. Amerman, Michael P. Cancro, Montserrat C. Anguera

**Author notes:** Correspondence: Montserrat C. Anguera.

## Abstract

Systemic lupus erythematosus (SLE) is an autoimmune disease preferentially observed in females. X-linked gene expression in XX females is normalized to that of XY males by X-Chromosome Inactivation (XCI). However, B cells from female SLE patients and mouse models of SLE exhibit mislocalization of Xist RNA, a critical regulator of XCI, and aberrant expression of X-linked genes, suggesting that impairment of XCI may contribute to disease. Here, we find that a subset of female mice harboring a conditional deletion of *Xis*t in B cells (“Xist cKO”) spontaneously develop SLE phenotypes, including expanded activated B cell subsets, disease-specific autoantibodies, and glomerulonephritis. Moreover, pristane-induced SLE-like disease is more severe in Xist cKO mice. Activated B cells from Xist cKO mice with SLE phenotypes have increased expression of proinflammatory X-linked genes implicated in SLE. Together, this work indicates that impaired XCI maintenance in B cells directly contributes to the female-bias of SLE.

## INTRODUCTION

Systemic lupus erythematosus (SLE) exhibits a strong female sex bias^1-3^ but the mechanisms underlying this sex bias are not well understood^2,4^. While there is some evidence for sex hormone contributions to this bias^5-7^, epidemiological studies demonstrate that SLE risk increases with the number of X chromosomes carried by an individual, suggesting important genetic contributions from the X chromosome. SLE risk in males with Klinefelter syndrome (47, XXY) is 14-fold higher than 46, XY males, and is similar to 46, XX females^8^. Further, the prevalence of SLE in females with polysomy X (XXX) is roughly twice that of 46, XX females, and more than twenty-fold that of 46, XY males^9^. Together, these observations indicate that X chromosome dosage contributes to SLE susceptibility.

To ensure dosage compensation of X-linked genes relative to XY males, mammals with multiple X chromosomes utilize X-chromosome inactivation (XCI), a complex and multi-layered epigenetic mechanism. XCI is initiated in early embryonic development when the long non-coding RNA Xist is upregulated from the future inactive X chromosome (Xi). Xist RNA then coats the Xi in *cis,* resulting in an allele-specific accumulation of multiple epigenetic modifications, including repressive histone marks (H3K27me3, H2AK119-ubiquitin), the histone variant macroH2A, and DNA methylation^10^. These heterochromatic modifications silence most genes across the Xi, except for some that homeostatically escape silencing and are expressed biallelically from the Xi and the active X chromosome (Xa)^11-14^. While the choice of which X to silence during XCI initiation is random, transcriptional repression of the chosen Xi is maintained throughout the lifetime of a cell and by its clonal daughters. The fidelity of XCI maintenance in female somatic cells is therefore an important regulator of X-linked gene dosage in XX individuals.

In contrast to other somatic cell types, B cells exhibit a unique mechanism of XCI maintenance^15^. The Xi in naive B cells lacks cytological enrichment of Xist RNA and some heterochromatic histone modifications, and antigen stimulation triggers the return of these modifications to the Xi^15,16^. However, circulating B cells isolated from female SLE patients exhibit features of impaired XCI maintenance, including abnormal XIST RNA localization patterns, reduced enrichment of heterochromatic H3K27me3 and H2AK119-ubiquitin modifications at the Xi, and aberrant X-linked gene expression^15,17^. Furthermore, lymphocytes from several mouse models of spontaneous lupus-like disease with varying degrees of female bias (NZB/W F1, NZM2328, and MRL/lpr) also have aberrant Xist RNA localization patterns and reduced heterochromatic mark enrichment at the Xi^18,19^. Given that B cells contribute to SLE pathogenesis^20-29^ and that the X chromosome is enriched for genes with immunomodulatory functions that are particularly relevant to B cell function, these findings have important implications for the female-biased development of SLE. Indeed, many X-linked genes with immune functions, including *TLR7*^30-32^, *TASL*^33^, *BTK*^34^, *IRAK1*^35^, *CD99*^36^, *IL9R*^37^ and *CXCR3*^38^, are overexpressed in peripheral blood lymphocytes from patients with SLE^39^. Moreover, *Tlr7*/*TLR7* gain-of-function in both mouse models and humans confers lupus-like phenotypes^32,40^. Together, these observations suggest that impaired XCI maintenance in B cells may result in the aberrant expression of immunomodulatory X-linked genes that contribute to the pathogenesis of SLE. However, it is not currently known whether disrupted XCI maintenance in B cells can cause SLE.

In this study, we asked whether perturbing XCI maintenance in B cells using a B-cell-specific *Xist* deletion could spontaneously cause SLE-like disease or enhance disease severity in a model of chemically-induced SLE. Using *Mb1*^Cre/WT^ *Xist*^fl^/^fl^ mice (Xist cKO), we discovered that some female Xist cKO mice spontaneously develop features of human SLE, including disease-specific anti-nuclear autoantibodies, proliferative glomerulonephritis, and increased numbers of activated B cells implicated in SLE pathogenesis, including age-associated B cells (ABCs)^41-44^ and plasma cells^45^. Furthermore, following pristane injection, female Xist cKO mice developed more severe SLE-like disease compared to their WT counterparts, with higher levels of anti-dsDNA autoantibodies and more activated B cell subsets including class-switched B cells, GL7+ activated B cells, plasma cells, and ABCs compared to controls. Finally, ABCs and GL7+ activated B cells from Xist cKO mice overexpressed the X-linked immunity gene *Tasl*, a member of the TLR7 signaling pathway that has been implicated in SLE pathogenesis^33,46,47^. Taken together, our work demonstrates that the disruption of XCI maintenance in B cells via the deletion of *Xist* can both spontaneously confer and exacerbate SLE-like disease, thereby providing a novel pathogenic mechanism that simultaneously accounts for the strong female sex bias of SLE.

## RESULTS

### Female Xist cKO mice on a nonautoimmune-prone background spontaneously develop features of human SLE

We used the *Mb1-Cre* transgene to delete *Xist* in the B cell lineage starting at the early pro-B cell stage, after the maintenance phase of XCI has been established (**Figure 1A**). *Xist*^2lox/2lox^ mice were successively mated to *Mb1*^WT/cre^ mice to derive heterozygous “Xist +/cKO” (*Xist*^WT/fl^; *Mb1*^WT/cre^) and homozygous “Xist cKO/cKO” (*Xist*^fl/fl^; *Mb1*^WT/cre^) mice. *Xist* deletion was confirmed in CD23+ splenic B cells using sequential Xist RNA fluorescence *in situ* hybridization (FISH) and immunofluorescence for H2AK119-ubiquitin (**Figures S1A-S1C**) and qPCR (**Figure S1D**), which demonstrated *Xist* deletion from the Xi in 100% of Xist cKO/cKO cells and ∼50% of Xist +/cKO cells, consistent with random XCI. Female Xist +/cKO and Xist cKO/cKO mice are viable with comparable survival to WT controls (**Figure S1E**). Adult female Xist cKO mice on a C57/BL6 background aged 6-8 months have similar percentages and numbers of total B220+ B cells compared to wildtype female mice in both the spleen (**Figure S1F)** and the bone marrow (**Figure S1G**), suggesting that *Xist* loss in the B cell compartment does not impact B cell development. Surprisingly, some female BALB/c Xist +/cKO mice (n=2 of 12) and Xist cKO/cKO mice (n=5 of 27) aged ≥12 months spontaneously developed elevated serum anti-dsDNA autoantibodies (**Figure 1B, Figure S2A-B**) at comparable levels to those observed in female NZB/W mice, a heavily female-biased murine model of spontaneous SLE-like disease^48^. These data indicate that the frequency of Xist cKO/cKO female mice spontaneously developing serologic features of SLE is around 18%. In a cohort of n=3 female Xist cKO/cKO mice with elevated serum anti-dsDNA (“Xist cKO High”), n=4 age-matched female Xist cKO/cKO mice with low serum anti-dsDNA (“Xist cKO Low”), and n=7 age-matched female WT mice (“WT”), we found that Xist cKO High mice not only have significantly higher serum levels of anti-dsDNA autoantibodies (**Figure 1C**) as measured by ELISA, but also significantly more anti-dsDNA autoantibody-secreting cells in both the spleen (**Figure 1D, Figure S2E**) and the bone marrow (**Figure S2C, Figure S2F**) as measured by anti-dsDNA ELISPOT. We also found that Xist cKO High mice had significantly higher levels of serum autoantibodies against Smith-U1RNP, a ribonuclear protein complex containing the Smith antigen, a highly SLE-specific antigen observed in some patients with SLE^49^ (**Figure 1E**).

**Figure 1.**
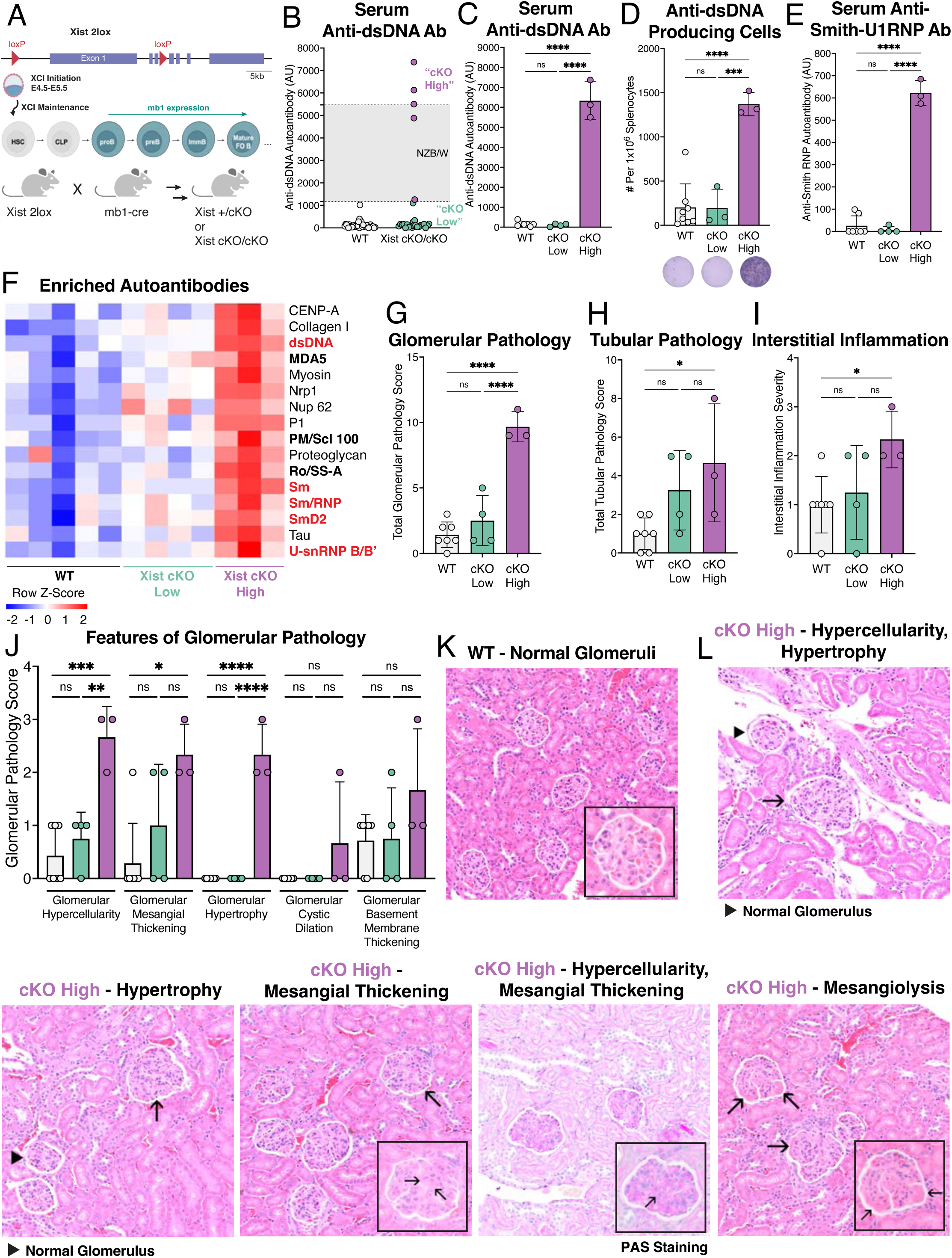
Female Xist cKO mice spontaneously develop features of SLE. **A.** Breeding schematic used to generate mice with a B cell-specific *Xist* deletion. Xist 2lox mice^85^ with loxP sites flanking the first 3 exons of *Xist* were successively bred to mb1-cre mice^86^ for deletion in the B cell lineage staring with early proB cells, to generate mice with a homozygous *Xist* deletion (“Xist cKO/cKO”) or a heterozygous *Xist* deletion (“Xist +/cKO”) in the B cell compartment. **B.** Serum levels of anti-dsDNA autoantibodies from female BALB/c mice 12 months or older (n=39 female WT mice, n=27 female Xist cKO/cKO mice) measured by enzyme-linked immunosorbent assay (ELISA). Each circle represents an individual mouse. Grey bar bounded by dotted lines represents serum levels of anti-dsDNA in NZB/W (n=8) mice that spontaneously develop SLE-like disease^48^. **C.** Mean serum anti-dsDNA autoantibody levels ± standard deviation (SD) in female Xist cKO/cKO mice with elevated anti-dsDNA autoantibodies (“Xist cKO High”, n=3), age-matched female Xist cKO/cKO mice with low anti-dsDNA autoantibodies (“Xist cKO Low”, n=4), and age-matched female WT controls (“WT”, n=7) as measured by ELISA. **D.** Mean number of anti-dsDNA producing cells per 1,000,000 splenocytes ± SD as measured by ELISPOT. **E.** Mean serum anti-Smith RNP autoantibody levels ± SD as measured by ELISA. **F.** Heatmap depicting individual sample-level data for those autoantibodies significantly enriched in Xist cKO High mice compared to WT mice or Xist cKO Low mice, as measured by an autoantigen microarray. Bold autoantibodies in black are either specific for or associated with non-SLE female biased autoimmune diseases. **G.** Mean total glomerular pathology ± SD. **H.** Mean total tubular pathology ± SD. **I.** Mean interstitial inflammation ± SD. **J.** Mean scores ± SD for individual features of glomerular pathology. **K.** Representative images of glomeruli from a WT mouse. **L.** Representative images of glomeruli from Xist cKO High mice.

Given the identification of multiple disease-specific autoantibodies in Xist cKO High mice, we sought to determine whether the sera of these mice contained additional autoantibodies implicated in autoimmunity. We performed autoantigen microarray profiling of serum from this cohort and found that Xist cKO High mice exhibited statistically significant enrichment of 16 of 120 autoantibodies examined (**Table S1**) compared to Xist cKO Low and WT samples. Xist cKO High serum contained significant enrichment of autoantibodies directed against the highly SLE-specific Smith antigens (Smith and SmithD2), the U1-RNP complex, and anti-dsDNA. Xist cKO High mice also had serum enrichment of autoantibodies associated with other female-biased autoimmune diseases including inflammatory myositis, systemic sclerosis, and Sjögren’s disease (MDA5, PM/Scl-100, and Ro/SS-A) (**Figure 1F**). Importantly, enrichment of these autoantibodies was not due to higher serum levels of total IgG in Xist cKO High mice (**Figure S2D**). While some autoantibodies from the array were decreased in Xist cKO High mice (**Figure S2G**), they displayed less consistent expression patterns across samples than the enriched autoantibodies. Furthermore, while Xist +/cKO mice tend to develop lower serum levels of anti-dsDNA compared to Xist cKO/cKO mice (**Figure S2A**), a rare female Xist +/cKO mouse with elevated anti-dsDNA also displayed enrichment of serum autoantibodies enriched in Xist cKO High mice (**Figure S2H**).

Given the spontaneous development of SLE-specific autoantibodies, we sought to determine whether Xist cKO High mice also developed renal pathology characteristic of lupus nephritis. Xist cKO High mice demonstrated pronounced features of focal (<50% of glomeruli affected) glomerular kidney disease (**Figure 1G**), recapitulating cardinal histopathologic manifestations of lupus nephritis. Xist cKO High mice also had modestly increased tubular disease (**Figure 1H**) and interstitial inflammation (**Figure 1I**) compared to controls. Xist cKO High mice developed variably moderate to sometimes severe glomerular hypercellularity, glomerular mesangial thickening, and glomerular basement membrane thickening with frequent secondary glomerular hypertrophy (**Figure 1J,L**). The glomerular hypercellularity in this subset of mice was specifically characterized by both segmental to global endocapillary loop hypercellularity and mesangial hypercellularity. Thus, about 18% of female Xist cKO mice over 12 months old with a B-cell specific *Xist* deletion spontaneously develop multiple autoantibodies associated with female-biased autoimmune diseases and glomerular pathology associated with SLE.

### Xist cKO High mice have increased numbers of activated B cell subsets associated with human SLE and murine SLE-like disease

Because we found that some Xist cKO/cKO mice spontaneously develop SLE pathologies, we next sought to identify changes in the B cell compartment that could contribute to these disease phenotypes. Xist cKO High mice had similar numbers of total splenic B cells and similar frequencies of both splenic and bone marrow B cells (**Figure S2I-J**) compared to Xist cKO Low and WT mice. However, an expansion in percentage and total cell number of several activated splenic B cell subsets were observed in Xist cKO High mice, including an expansion of CD11c+ age-associated B cells (“ABCs”; B220+CD19+CD11c+) (**Figure 2B**), which are elevated in SLE patients and mouse models of lupus-like disease^50,51^ and splenic GL7+ B cells (B220+CD19+IgM-IgD-GL7+), an activated population that includes germinal center B cells, which are expanded in mouse models of SLE^52^ (**Figure 2C**). Xist cKO mice had an increased number of IgM-IgD- class-switched B cells compared to controls (**Figure 2D**). Xist cKO High mice also exhibited increased numbers of splenic short-lived plasma cells (“SLPCs”; IgD-B220+CD138+), considered to be the primary producers of anti-dsDNA autoantibodies in SLE^45^, and long-lived plasma cells (“LLPCs”; IgD-B220-CD138+) were modestly increased (**Figure 2E**). Interestingly, 6-month-old C57/BL6 Xist cKO/cKO mice demonstrate some increases in activated splenic B cell subsets, including a slightly higher number of GL7+ B cells (**Figure S2K**), class switched B cells (**Figure S2L**), and short-lived plasma cells (**Figure S2M**), but not CD11c+ ABCs (**Figure S2N**) or long-lived plasma cells (**Figure S2O**), suggesting that some of these populations may expand before the development of serologic autoimmunity. In contrast to the splenic plasma cell pool, the bone marrow plasma cell pool is not expanded in Xist cKO High mice compared to Xist cKO Low and WT mice (**Figure S2P**) despite our finding that Xist cKO High mice have more anti-dsDNA producing cells in the bone marrow compared to controls (**Figure S2C**). In summary, these findings demonstrate that loss of *Xist* in the B cell compartment leads to spontaneous development of serologic, histopathological, and cellular features of human SLE in a non-autoimmune mouse background, suggesting that appropriate maintenance of XCI in B cells is critical for restricting autoreactive immune processes.

**Figure 2.**
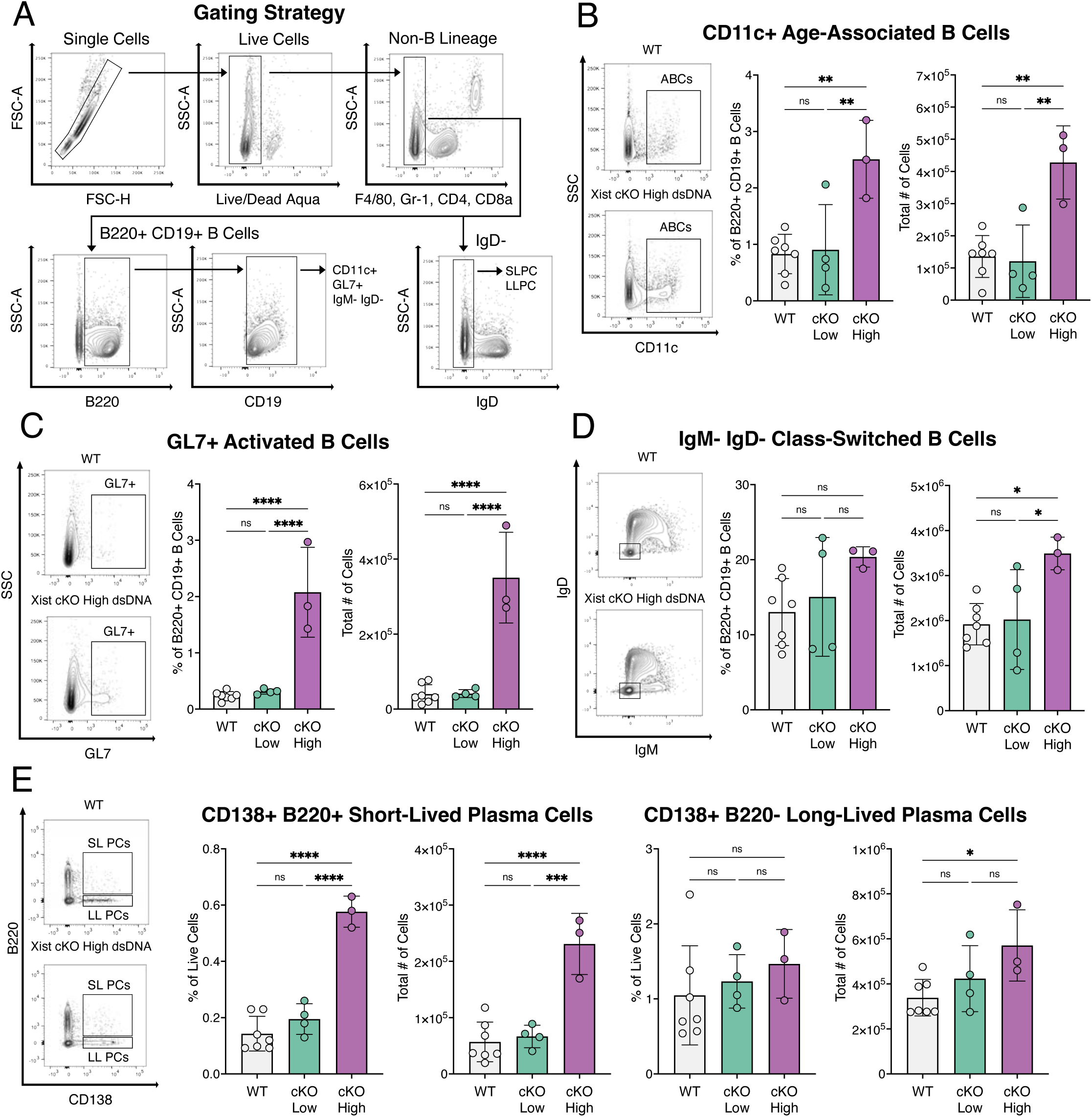
Activated B cell subsets expand in Xist cKO High mice with serologic and pathologic features of SLE. **A.** Gating strategy for flow cytometry analysis of splenocytes. **B-E.** Cells were pre-gated on live single cells that are CD4-CD8-Gr-1-F4/80-. Quantification of splenic (**B**) age-associated B cells (ABCs; B220+CD19+CD11c+), (**C**) GL7+ activated B cells (B220+CD19+IgM-IgD-CD138-GL7+), (**D**) class-switched B cells (B220+CD19+IgM-IgD-), and (**E**) short-lived plasma cells (SLPCs; IgD-CD138+B220+) and long-lived plasma cells (LLPCs; IgD-CD138+B220-). Data are depicted as mean ± SD. * *p* < 0.05, ** *p* < 0.01, *** *p* < 0.001, **** *p* < 0.0001 via ordinary one-way ANOVA with Tukey’s multiple comparisons test.

### Age-associated B cells from Xist cKO mice with spontaneous SLE-like pathology upregulate expression of the TLR7 pathway member Tasl

To determine how *Xist* loss affects X-linked gene expression and contributes to the development of SLE-like disease, we performed RNA sequencing on B cell subsets from Xist cKO High (n=3, 17.9 months ± 1.3), Xist cKO Low mice (n=4, 17.9 months ± 4.0) and WT controls (n=6, 18.9 months ± 3.4). First, we sequenced splenic CD23+ B cells, which include mature naive follicular B cells that serve as progenitors for activated B cell subsets including germinal center, memory and plasma cells^53^. Principal component analyses (PCA) distinguished CD23+ B cells from Xist cKO High mice, while Xist cKO Low and WT mice were indistinguishable (**Figure S3A**). We observed 2-3 X-linked genes that were differentially expressed in both Xist cKO High (**Figure 3A-B**) and Xist cKO Low (**Figure 3C-D**) CD23+ B cells relative to WT controls (**Table S2-3**), which includes the expected down-regulation of *Xist* in Xist cKO samples (**Figure S3B**). The small number of differentially expressed X-linked genes may reflect the cellular heterogeneity of the CD23+ B cell pool in Xist cKO High mice, which may mask changes in X-linked gene expression occurring in individual CD23+ cellular subsets. Notably, CD23+ B cells from Xist cKO High, but not Xist cKO mice, exhibited significant upregulation of X-linked *Kif4* (kinesin family member 4 protein) (**Figure 3A-B**). Examination of autosomal transcriptional changes between Xist cKO High and WT mice reveals altered expression of n=86 genes (**Figure 3E, Figure S3C, Table S3**), and upregulated DEGs are enriched for genes implicated in cell cycle, cell division, and proliferation (**Table S9**), likely reflecting the expansion of GL7+ activated CD23+ cells in Xist cKO High mice (**Figure S3D**).

**Figure 3.**
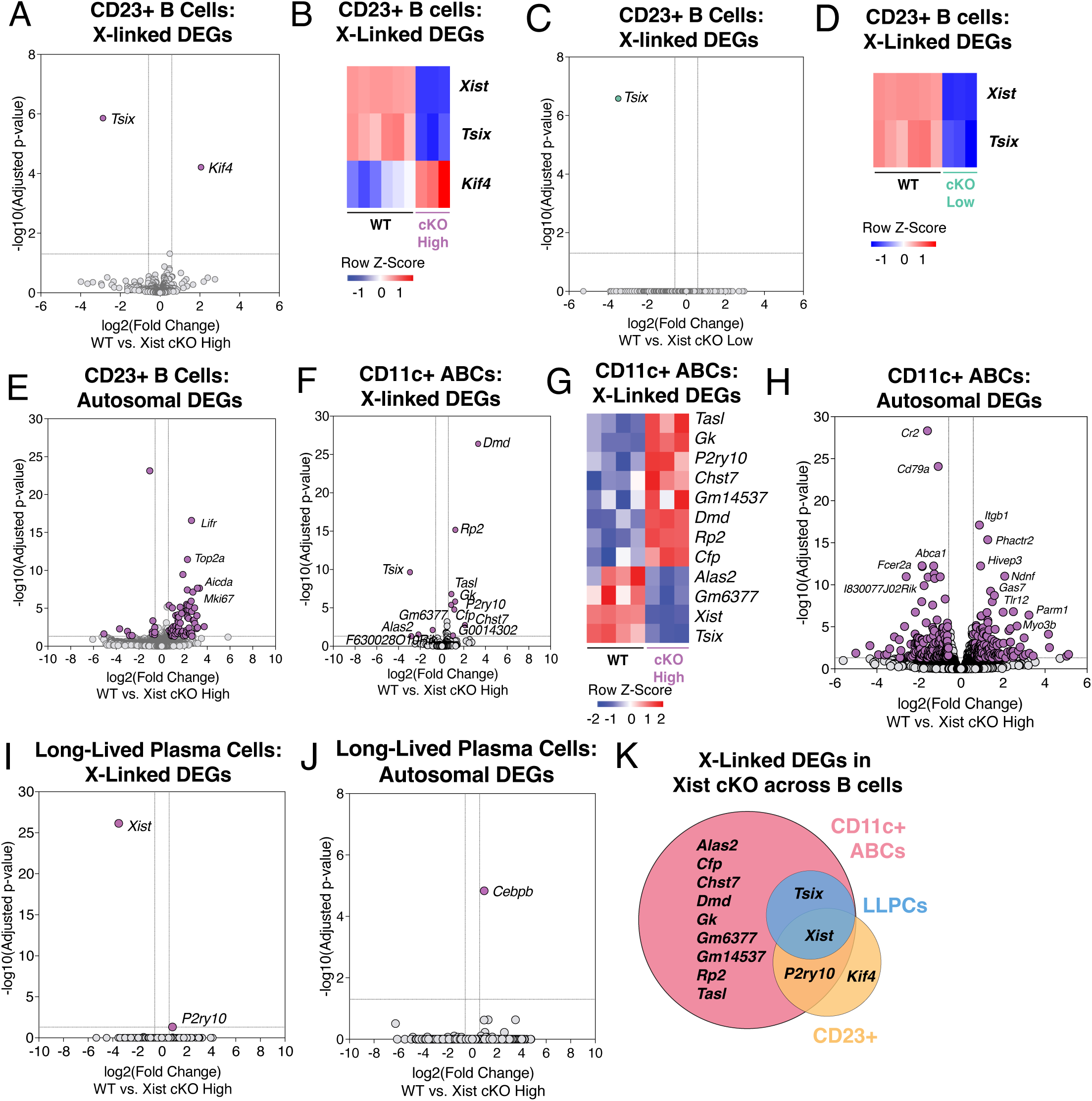
X-linked *Tasl* is upregulated in age-associated B cells from Xist cKO High mice with SLE-like disease. **A.** Volcano plot of X-linked genes that are significantly differentially expressed (purple) between Xist cKO High and WT splenic CD23+ B cells. **B.** Heatmap of differentially expressed X-linked genes between Xist cKO High and WT splenic CD23+ B cells. **C.** Volcano plot of X-linked genes that are significantly differentially expressed (green) between Xist cKO Low and WT splenic CD23+ B cells. **D.** Heatmap of differentially expressed X-linked genes between Xist cKO Low and WT splenic CD23+ B cells. **E.** Volcano plot of autosomal genes that are significantly differentially expressed (purple) in CD23+ B cells from Xist cKO High and WT mice. **F.** Volcano plot depicting X-linked genes that are significantly differentially expressed (purple) between Xist cKO High and WT CD11c+ ABCs. **G**. Heatmap of differentially expressed X-linked genes between Xist cKO High and WT CD11c+ ABCs. **H.** Volcano plot of autosomal genes that are significantly differentially expressed (purple) in CD11c+ ABCs from Xist cKO High and WT mice. **I.** Volcano plot depicting X-linked genes that are significantly differentially expressed (purple) between Xist cKO High and WT LLPCs. **J.** Volcano plot of autosomal genes that are significantly differentially expressed (purple) in SLPCs from Xist cKO High and WT mice. **K.** Overlap of X-linked DEGs from CD11c+ ABCs (pink), LLPCs (blue), and CD23+ B cells (yellow) from Xist cKO mice.

We next examined transcriptional changes in splenic B cell subsets that are expanded in Xist cKO High mice relative to Xist cKO Low mice or WT mice: CD11c+ ABCs and CD138+ B220- long-lived plasma cells (LLPC). TLR7 (X-linked) drives ABC activation and differentiation^54^, suggesting that this cell type is likely to be particularly sensitive to gene expression changes of X-linked TLR7 pathway members (*Tlr7, Tasl*, and *Irak1*). Gene expression profiles of ABCs from Xist cKO High (n=3) and WT (n=4) mice were distinct by PCA (**Figure S3E**). We observed 8 upregulated X-linked genes in Xist cKO High ABCs (**Figure 3F-G, Table S4**), and only 4 downregulated X-linked genes including *Xist* (**Figure S3F**). Xist cKO High ABCs have increased expression of *Tasl,* the aforementioned X-linked TLR7 pathway member. *TASL* (*CXORF21*) was first identified by GWAS as a potential causal variant of SLE^33^, and is upregulated in immune cells from SLE patients. TASL is particularly important for the signal transduction cascade downstream of endosomal TLRs including TLR7^46,47^. Expression of *Tlr7* was slightly, but not statistically significantly increased in Xist cKO ABCs compared to WT (**Figure S3G**) suggesting that multiple members of the TLR7 pathway may be upregulated in Xist cKO High mice to varying degrees. More generally, there is aberrant transcription genome-wide in ABCs from Xist cKO High mice, where 142 genes are upregulated and 166 are downregulated (**Figure 3H, Figure S3H, Table S4**). Differentially upregulated genes are enriched for pathways including lymphocyte activation, inflammatory response, and lymphocyte proliferation (**Table S9**), which may contribute to CD11c+ ABC cellular expansion in Xist cKO mice. Downregulated genes were also enriched for genes involved in immune response pathways (**Table S9**). Unlike ABCs, Xist cKO High LLPCs do not segregate from WT LLPCs based on PCA (**Figure S3I**), and differential gene expression analysis revealed minimal X-linked (**Figure 3I**) and autosomal (**Figure 3J**) gene expression differences, suggesting that LLPCs are not transcriptionally sensitive to *Xist* loss. In summary, these findings indicate that CD11c+ ABCs, which expand in Xist cKO High mice, are particularly sensitive to *Xist* deletion, as they exhibit the greatest numbers of differentially expressed X-linked genes compared to LLPCs and CD23+ B cells (**Figure 3K**). In particular, *Xist* deletion in the B cell compartment results in the increased expression of *Tasl* within ABCs, but not LLPCs or CD23+ B cells. This suggests that *Tasl* overexpression due to impaired XCI maintenance in ABCs may promote their expansion and ultimately, their role in the spontaneous development of SLE-like disease in a female-biased manner.

### Pristane treated Xist cKO/cKO mice develop more severe SLE-like disease

Spontaneous lupus-like disease is not fully penetrant in female Xist cKO mice. Thus, to determine how X-linked dosage imbalances impact B cell function in the context of autoimmunity, we used pristane to induce SLE-like disease in female Xist cKO mice and WT littermates. Pristane-induced SLE-like disease exhibits a female bias^55^, and recapitulates a number of features of human disease including elevated nuclear autoantibodies (anti-dsDNA, chromatin, Sm, RNP), glomerulonephritis^56^, and the Type 1 interferon-stimulated gene signature observed in the peripheral blood of patients with SLE^57^. Furthermore, the development of pristane-induced disease requires X-linked *Tlr7* signaling, the MyD88 adapter protein, and the Type I IFN receptor^58-61^, suggesting that this model relies on molecular drivers similar to those implicated in human SLE^32^. We therefore injected female BALB/c Xist cKO (n=36) and WT (n=46) mice with pristane or WT mice with PBS (n=10) as a negative control. We monitored proteinuria and serum anti-dsDNA levels within these cohorts over 9 months and collected spleens for flow cytometry at 4, 7, or 9 months post pristane injection and kidneys for pathological assessment at 7 months post pristane injection (**Figure 4A**). All pristane-treated mice developed elevated levels of anti-dsDNA autoantibodies starting at 3 months post-injection, and Xist cKO mice developed significantly higher serum anti-dsDNA levels compared to pristane-treated female WT mice over 7 months of disease (**Figure 4B**). Pristane-treated Xist cKO mice also had higher levels of serum anti-dsDNA at 9 months of disease (**Figure 4C**). We did not observe differences in proteinuria between Xist cKO and WT mice over 9 months of disease (**Figure S4A**). However, pristane-treated female Xist cKO mice had slightly more severe total glomerular kidney pathology (**Figure 4D**) and similar tubular kidney pathology (**Figure 4E**) and interstitial inflammation (**Figure 4F**) compared to WT mice at 7 months post-injection. Specifically, Xist cKO mice treated with pristane demonstrated relatively more pronounced glomerular hypercellularity, as well as glomerular mesangial thickening, basement membrane loop thickening, and secondary glomerular hypertrophy compared to WT pristane-treated mice (**Figure 4G**). These mice also developed several other morphologic changes within glomeruli that are associated with active glomerulonephritis in humans with lupus nephritis^62^, including crescent formation and mesangiolysis (**Figure 4J**). Thus, Xist cKO mice with pristane-induced SLE-like disease developed more severe serologic autoimmunity and slightly more severe glomerular pathology compared to pristane-treated WT mice.

**Figure 4.**
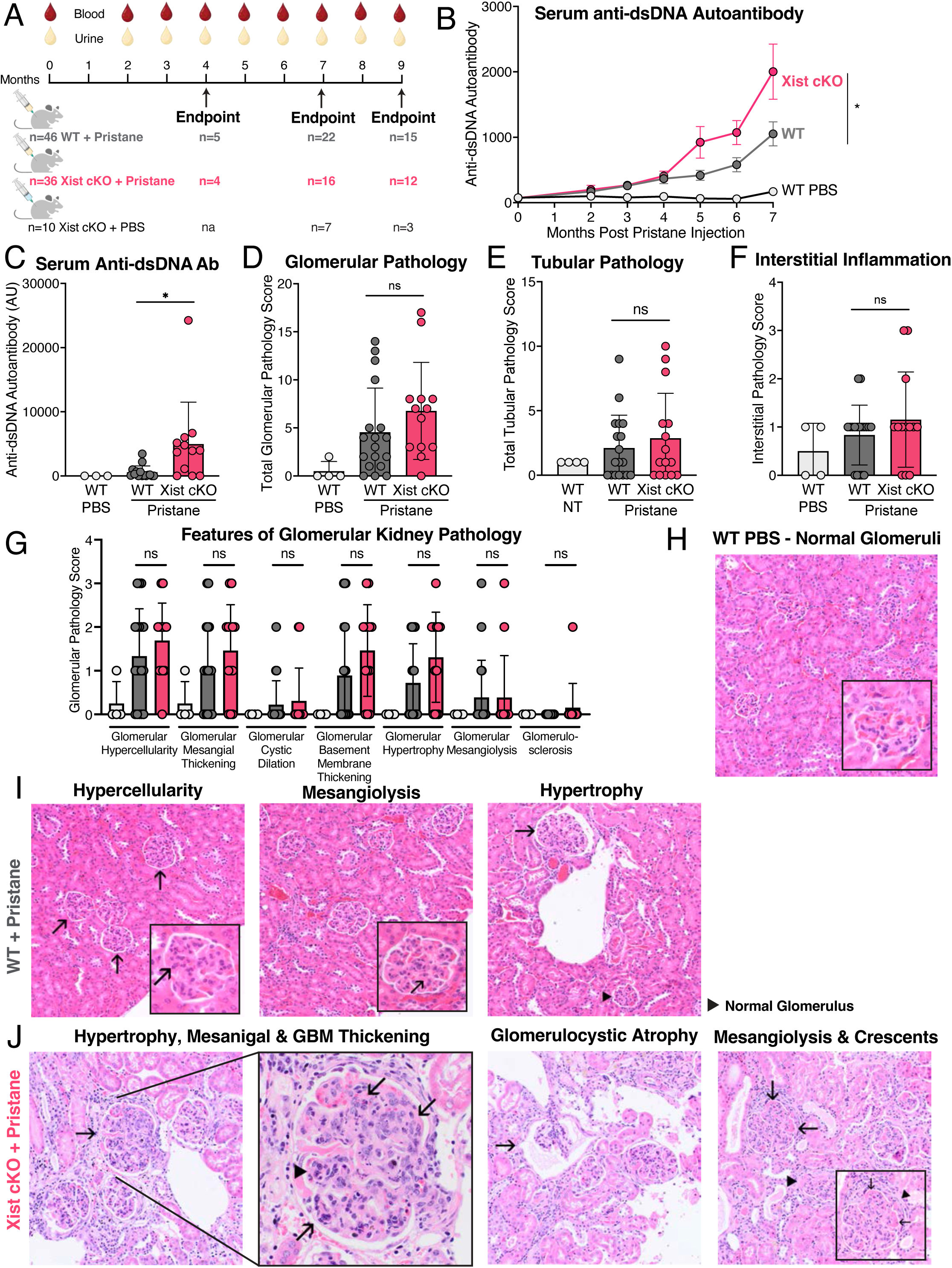
Xist cKO mice develop more severe features of pristane-induced SLE. **A.** Experimental design for inducing SLE-like disease with pristane. Female WT and Xist cKO mice were injected with pristane and monitored for up to 9 months for development of elevated autoantibodies in the blood and proteinuria. At experimental endpoint, organs were harvested for flow cytometry and pathologic assessment. N=4 WT mice and n=4 Xist cKO mice died spontaneously during disease development, and are not included in Endpoint cohorts. **B.** Mean serum anti-dsDNA autoantibody levels ± SEM in pristane-treated Xist cKO mice (n=32), pristane-treated WT mice (n=41), and PBS-treated WT mice (n=10) over 7 months. *p* < 0.05 by mixed effects model with the Geisser-Greenhouse correction (not assuming equal variability of differences). **C.** Mean serum anti-dsDNA autoantibody levels ± SD in pristane-treated Xist cKO mice (n=12), pristane-treated WT mice (n=18), and PBS-treated WT mice (n=3) at 9 months post-pristane injection. **D.** Mean total glomerular pathology score ± SD. **E.** Mean total tubular pathology score ± SD. **F.** Mean interstitial inflammation ± SD. **G.** Mean scores for individual features of glomerular pathology ± SD. * *p* < 0.05 via pairwise Mann-Whitney U test comparing pristane-treated Xist cKO to pristane-treated WT mice. **H.** Representative image of glomeruli from a WT-PBS mouse. **I**. Representative image of glomeruli from pristane-treated WT mice. **J**. Representative image of glomeruli from pristane-treated Xist cKO mice.

Next, we examined the impact of a B-cell-specific *Xist* deletion on splenic B cell subsets in the context of pristane-induced SLE-like disease at 4, 7, and 9 months post-pristane injection. Pristane-treated Xist cKO mice had a similar frequency and number of splenic B220+ B cells at all timepoints measured compared to pristane-treated WT mice, and the frequency of B cells contracted in pristane-treated mice compared to non-treated controls after 9 months of disease (**Figure S4B**). Pristane-treated Xist cKO mice had a higher frequency of class-switched splenic B cells compared to pristane-treated WT mice at 4, 7, and 9 months post-pristane injection; however, the total number of class-switched splenic B cells was greater in Xist cKO pristane mice only at 9 months post-pristane (**Figure 5A**). GL7+ activated B cells, which include germinal center B cells, were enriched in frequency in Xist cKO pristane mice at all timepoints measured, and in number at 7 and 9 months post-pristane (**Figure 5B**). Short-lived plasma cells were enriched in frequency and number in Xist cKO pristane mice at 7- and 9-months post pristane (**Figure 5C**), while long-lived plasma cells were increased in frequency in Xist cKO mice at 9 months post pristane (**Figure S4C**). Finally, CD11c+ ABCs were enriched in Xist cKO mice in frequency at 7 and 9 months post-pristane and in number at 9 months post pristane compared to pristane-treated WT mice (**Figure 5D**). These findings indicate a temporal distinction in the expansion of activated B cell subsets in pristane-treated Xist cKO mice, with expansion of class-switched and GL7+ B cells as early as 4 months post-pristane, followed by expansion of short-lived plasma cells and CD11c+ ABCs at 7 months post-pristane. Additionally, these data demonstrate that *Xist* deletion in B cells leads to expansion of activated B cell subsets during both spontaneous and pristane-induced SLE. Thus, *Xist* deletion in B cells alters the B cell compartment and results in the gradual accumulation of activated splenic B cell subsets implicated in response to pristane, ultimately resulting in more severe SLE-like disease.

**Figure 5.**
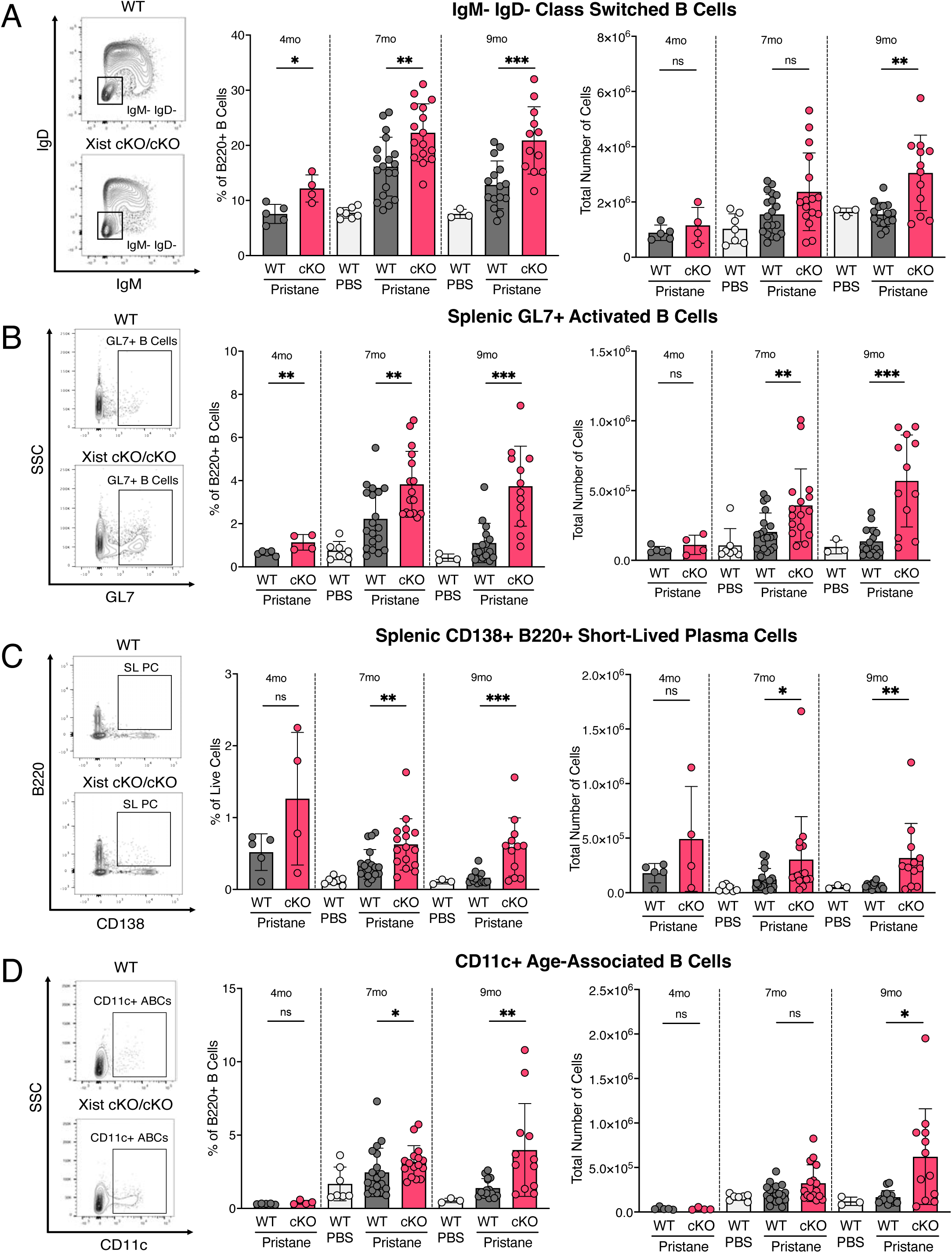
Pristane-treated Xist cKO mice develop more activated B cells compared to pristane-treated WT mice. **A-D.** Cells were pre-gated on live, single CD4-CD8-Gr-1-F4/80-cells. Quantification of splenic (**A**) class-switched B cells (B220+CD19+IgM-IgD-), (**B**) GL7+ activated B cells (B220+CD19+IgM-IgD-CD138-GL7+), (**C**) short-lived plasma cells (SLPCs; IgD-CD138+B220+), and (**D**) age- associated B cells (ABCs; B220+CD19+CD11c+) after 4, 7, or 9 months of pristane. Data are depicted as mean ± SD. * *p* < 0.05, ** *p* < 0.01 via pairwise Mann-Whitney U test comparing pristane-treated Xist cKO to pristane-treated WT mice.

### Xist +/cKO mice develop intermediate pristane-induced SLE phenotypes

Because some Xist +/cKO mice spontaneously developed SLE-like disease but with lower serum anti-dsDNA levels compared to Xist cKO/cKO mice (**Figure S2A**), we investigated the severity of pristane-induced SLE phenotypes in Xist +/cKO mice. We injected female BALB/c Xist +/cKO mice (n=23) and WT mice (n=46) and monitored animals for proteinuria and serum dsDNA autoantibodies over 9 months, and then examined kidney sections for pathological assessment and spleen for B cell subset analyses using flow cytometry (**Figure S5A**). Xist +/cKO mice developed similar levels of serum anti-dsDNA autoantibodies compared to pristane-treated WT mice (**Figure S5B**) over 7 months, yet had higher anti-dsDNA autoantibodies at 9 months post-pristane compared to WT pristane-treated mice (**Figure S5C**). Xist +/cKO mice developed similar levels of proteinuria over time compared to pristane-treated WT mice (**Figure S5D**). Glomerular and tubular pathology and interstitial inflammation were also similar between WT and Xist +/cKO mice after 7 months of pristane (**Figures S5E-G**). Heterozygous *Xist* deletion did not affect total B cell numbers, as the total splenic B cell pool was similar in WT and Xist +/cKO mice treated with pristane (**Figure S5H**). However, we observed changes in the splenic B cell compartment similar to but more subtle than those observed in Xist cKO/cKO mice. Expansions of activated B cells in pristane-treated Xist +/cKO mice, including class-switched B cells (**Figure S5I**), GL7+ activated B cells (**Figure S5J**), short-lived (**Figure S5K**) and long-lived plasma cells (**Figure S5L**), and CD11c+ ABCs (**Figure S5M**) were more modest compared to pristane-treated Xist cKO/cKO mice, and some expansions were delayed compared to mice with homozygous loss of *Xist*. These findings suggest that *Xist* deletion in approximately half of the B cell pool, as found in Xist +/cKO mice, yields milder pristane-induced phenotypes compared to Xist cKO/KO mice that have perturbed XCI in all of their B cells.

### Age-associated B cells and GL7+ activated B cells from pristane-treated Xist cKO mice upregulate Tasl

To determine how *Xist* loss impacts X-linked dosage compensation and contributes to the increased severity of pristane-induced disease observed in Xist cKO/cKO mice, we performed RNA sequencing on splenic B cell subsets expanded in Xist cKO mice treated with pristane for 7 months (age 10-12 months). ABCs from pristane-treated Xist cKO (n=4 mice) and WT (n=6 mice) were mostly transcriptionally distinct by PCA (**Figure S6A**) and demonstrated changes in X-linked gene expression (n=13 X-linked DEGs) (**Figure 6A-B**). Interestingly, we again observed that *Tasl* is upregulated in ABCs of Xist cKO mice treated with pristane, along with *Dmd, Rp2, Cfp*, and *Ftx*, which is a positive activator of *Xist* expression^63^. Of the 12 X-linked DEGs found in ABCs from Xist cKO mice with spontaneous SLE (**Figure 3F-G**) and the 13 X-linked DEGs found in ABCs from Xist cKO mice with pristane-induced SLE, 6 X-linked DEGs were shared, including downregulated genes *Xist* and *Tsix*, and upregulated genes *Cfp*, *Dmd*, *Rp2*, and *Tasl* (**Figure 6C**). We also observe that 264 autosomal genes were differentially expressed in Xist cKO ABCs (**Figure 6D, Figure S6B**), and that most genes (234 genes, 87% of all DEGs) were downregulated in pristane-treated Xist cKO ABCs compared to pristane-treated WT ABCs. Differentially downregulated genes were implicated in pathways of cytokine production, responses to external stimuli, and inflammatory responses (**Table S9**); differentially upregulated genes were enriched for immune response pathways and cellular pH pathways (**Table S9**).

**Figure 6.**
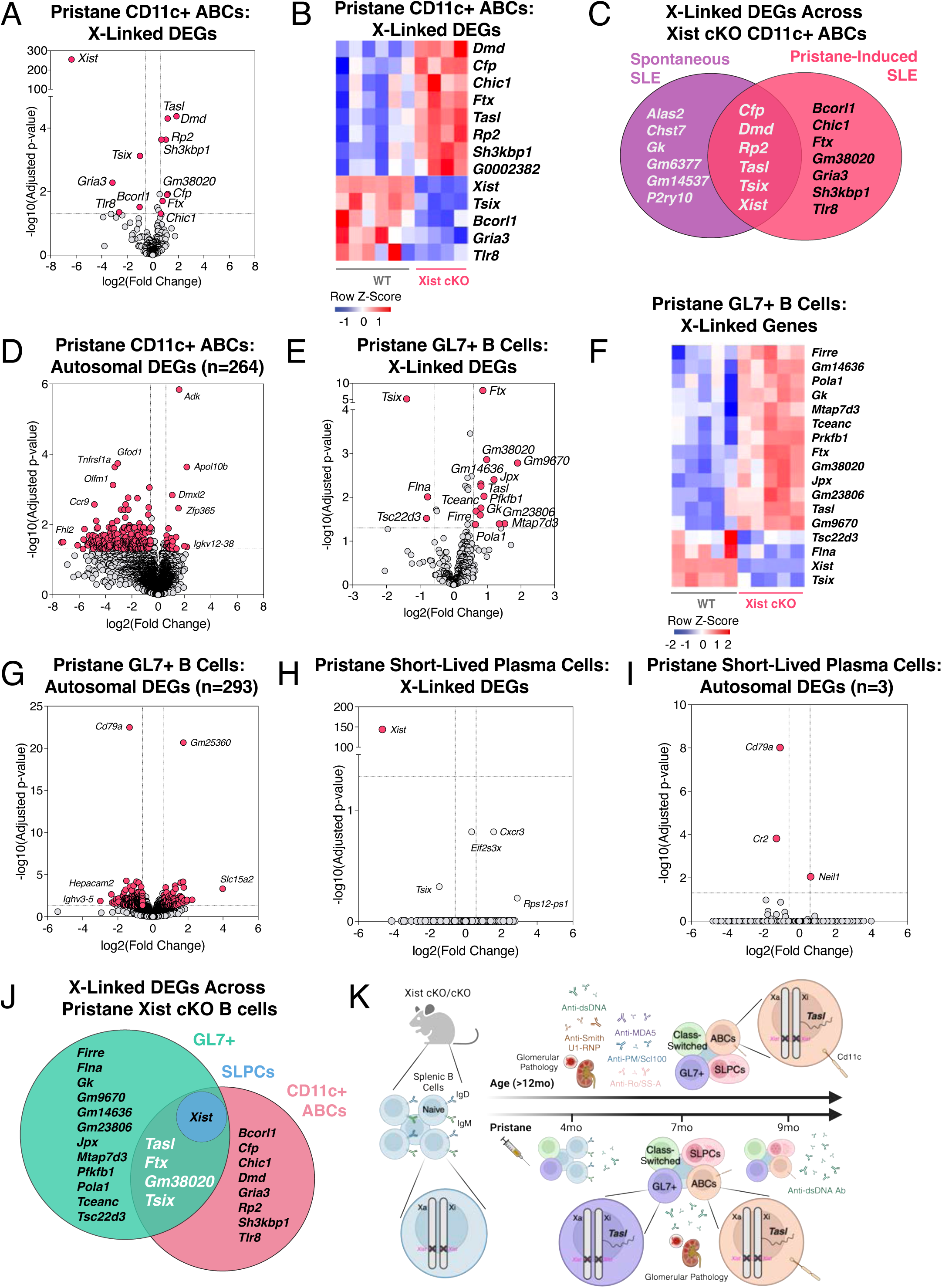
*Tasl* is upregulated in ABCs and GL7+ activated B cells from pristane-treated Xist cKO mice. **A.** Volcano plot depicting X-linked genes that are significantly differentially expressed (pink) in CD11c+ ABCs from pristane-treated Xist cKO vs. pristane-treated WT mice. **B.** Heatmap of differentially expressed X-linked genes in CD11c+ ABCs from pristane-treated mice. **C.** Overlap of X-linked DEGs found in CD11c+ ABCs from Xist cKO mice with spontaneous SLE (purple) and CD11c+ ABCs from Xist cKO mice with pristane-induced SLE (pink). **D.** Volcano plot depicting autosomal genes that are significantly differentially expressed (pink) in CD11c+ ABCs from pristane-treated Xist cKO vs. pristane-treated WT mice. **E.** Volcano plot depicting X-linked genes that are significantly differentially expressed (pink) in GL7+ activated B cells from pristane-treated Xist cKO vs. pristane-treated WT mice. **F**. Heatmap of differentially expressed X-linked genes in GL7+ B cells from pristane-treated mice. **G.** Volcano plot depicting autosomal genes that are significantly differentially expressed (pink) in GL7+ B cells from pristane-treated Xist cKO vs. pristane-treated WT mice. **H.** Volcano plot depicting X-linked genes that are significantly differentially expressed (pink) in SLPCs from pristane-treated Xist cKO vs. pristane-treated WT mice. **I**. Heatmap of differentially expressed X-linked genes in SLPCs from pristane-treated mice. **J.** Overlap of X-linked DEGs found among B cell subsets from pristane-treated Xist cKO mice. GL7+ B cells (green), SLPCs (blue), and CD11c+ ABCs (pink). **K.** Working model schematic depicting serologic, cellcular, pathologic, and X-linked transcriptional changes in Xist cKO mice with spontaneous SLE phenotypes (top) or pristane-induced SLE-like disease (bottom).

Splenic activated GL7+ B cells from pristane-treated Xist cKO (n=5 mice) and WT (n=5 mice) were also transcriptionally distinct by PCA (**Figure S6C**) and exhibit a significant number (n=17) of differentially expressed X-linked genes, of which the majority are upregulated (**Figure 6E-F**). Interestingly, among the upregulated X-linked genes in GL7+ B cells from Xist cKO mice, we again observe *Tasl* and *Ftx* (**Figure 6F**), which were upregulated in CD11c+ ABCs. Upregulated X-linked genes also include *Firre*, a long noncoding RNA that regulates 3D structure of the Xi and has a role in hematopoiesis, where its overexpression results in increased production of pro-inflammatory cytokines^64,65^. *Xist* was significantly downregulated, as expected (**Figure S6D**).

GL7+ B cells from pristane-treated Xist cKO mice exhibit differential gene expression of 293 autosomal DEGs (**Figure 6G, Figure S6E**); upregulated genes were associated with pathways involved in metabolism, while downregulated genes were associated with pathways involving cell activation, developmental processes, and tissue morphogenesis (**Table S9**). SLPCs from pristane-treated Xist cKO (n=6 mice) and WT (n=6 mice) were transcriptionally similar to each other (**Figure S6F**) and did not exhibit any differentially expressed X-linked genes beyond *Xist* downregulation (**Figure 6H**). Similarly, there were very few differentially expressed autosomal genes (**Figure 6I**). Comparison of X-linked DEGs across B cell subsets from pristane-treated Xist cKO female mice reveals mostly cell type-specific profiles, yet there are 3 X-linked genes upregulated in both CD11c+ ABCs and GL7+ activated B cells: *Tasl, Ftx, and Gm38020,* which overlaps with the *Ftx* gene (**Figure 6J**). In sum, *Xist* deletion in B cells alters the expression of X-linked genes in GL7+ B cells and CD11c+ B cells, with *Tasl* and *Ftx* being significantly upregulated. Taken together, our data demonstrate that perturbed XCI maintenance via *Xist* deletion in the B cell compartment results in the upregulation of X-linked *Tasl*, a contributor to abnormal immune signaling in SLE, along with activated B cell subset expansion, production of disease-specific autoantibodies, and glomerulonephritis. (**Figure 6K**).

## DISCUSSION

Here we demonstrate that *Xist* deletion in B cells of female BALB/c mice can result in the spontaneous development of SLE-like disease phenotypes, with highly specific features of SLE including serologic autoimmunity, expansion of disease-relevant B cell subsets, and glomerular-predominant kidney pathology. Furthermore, using pristane to chemically induce SLE-like disease, we show that female Xist cKO mice develop a greater expansion of specific B cell subsets and higher anti-dsDNA autoantibody levels compared to pristane-treated WT mice as SLE-like disease develops over time. In both spontaneous and chemically-induced SLE-like disease, *Xist* deletion in B cells alters X-linked gene expression in activated and/or memory B cells, and results in the upregulation of *Tasl*, a Tlr7-9 pathway member, suggesting a key role for this X-linked protein in driving altered B cell function, autoantibody production, and renal pathology in a sex-biased manner. Together, our data support the hypothesis that disrupted XCI maintenance, as observed in B cells from patients with SLE and female-biased murine models of SLE-like disease^15,17,19^, drives sex-biased systemic autoimmunity by modifying the dosage of proinflammatory X-linked genes.

Our results are in agreement with a recent publication demonstrating that global reduction of Xist expression following whole-body deletion of X-linked *Ftx*, a positive activator of *Xist* expression^66^, results in development of lupus-like disease. Although Xist RNA levels are only reduced by female Ftx KO mice by ∼50%, these mice exhibit upregulation of X-linked immunity genes including *Tlr7* in splenic B cells, dendritic cells, and monocyte/macrophages and *Tasl* in dendritic cells and monocyte/macrophages. Female Ftx KO mice also exhibit a higher frequency of myeloid and plasmacytoid dendritic cells, an expansion of activated splenic B and T cells, and development of SLE-specific serum autoantibodies. Thus, while we find that *Xist* deletion that is restricted to the B cell compartment is sufficient to induce spontaneous SLE, it remains possible that decreasing its expression in other cell types, as occurs the Ftx KO mouse model, will increase the frequency or severity of disease. Like our B-cell-specific Xist cKO mice, Ftx KO mice fail to develop some SLE disease features until one year of age. This commonality suggests that additional age-related epigenetic changes (either directly or indirectly) potentiate the effects of *Xist* loss.

Female Xist cKO mice on a BALB/c background spontaneously develop SLE pathologies, yet the frequency of disease is ∼18%, and only occurs in mice 12 months or older. While human SLE similarly exhibits a low penetrance within affected families, the average age for a woman to be diagnosed with SLE is 31 years, underscoring the contribution of additional factors beyond DNA sequence and age in the pathogenesis of SLE^67^. Indeed, a variety of environmental factors, including sex hormones, cigarette smoke, viral infections (such as Epstein-Barr virus), medications, silica exposure, dietary factors, and pollution/occupational chemicals, have been associated with the development of SLE^68,69^. In humans, exposure to these factors may drive epigenetic changes that could contribute to erosion of epigenetic silencing of the Xi in immune cells. As such, we posit that *Xist* deletion in female mice represents an underlying genetic/epigenetic susceptibility factor for SLE-like disease and that enivornmental or age-related epigenetic changes^70^ may also be required for the aberrant escape of X-linked genes and the subsequent development of SLE phenotypes. Another possibility is that an immune-activating event is required to trigger SLE in Xist cKO mice. Additional research is necessary to determine whether pathogens may accelerate the development of SLE-like disease in Xist cKO mice. Notably, we also find that Xist cKO High mice exhibit serologic features of other female-biased autoimmune diseases, including enrichment of autoantibodies against MDA5, PM/Scl-100, and Ro/SS-A. This may point to a role for disrupted XCI maintenance in B cells contributing to multiple female-biased autoimmune diseases, including Sjögren’s disease, systemic sclerosis, and amyopathic dermatomyositis.

*Xist* deletion in B cells leads to upregulation of numerous X-linked genes in spontaneous and pristane-induced SLE, with *Tasl* being the most consistently upregulated X-linked gene across B cell subsets. *Tasl* is an endosomal adaptor protein that functions in Tlr7 and Tlr9 signaling^47^ and is notably upregulated in peripheral blood mononuclear cells from patients with SLE^46^. Our finding is consistent with *in vitro* work demonstrating that *XIST* deletion in cultured human B cells leads to the upregulation of *TASL* as well as expansion of CD11c+ B cells^50^. Despite *Tlr7* being only slightly upregulated in our findings, *Tasl* upregulation may enhance signaling through the Tlr7 pathway, contributing to the expansion of activated B cell populations. Indeed, a gain-of-function mutation in *Tlr7* leads to an expansion of ABCs, germinal center B cells, and plasma cells^32^. Other X-linked genes whose genomic loci are proximal to the locus of *Tasl* were also upregulated in Xist cKO mice, including *Dmd* and *Gk*. Upregulation of these X-linked gene clusters in the context of *Xist* loss may suggest local changes in epigenetic modifications or genomic architecture on the Xi. Notably, *DMD* expression was also increased in cultured human B cells after CRISPR-Cas9 mediated deletion of *XIST*^71^. Intriguingly, *Ftx*, a positive regulator of *Xist* expression, was upregulated in both CD11c+ ABCs and GL7+ B cells in pristane-treated Xist cKO mice, suggesting a positive feedback loop that is induced during disease to upregulate *Xist* expression. Finally, *Firre*, which regulates 3D structure of the Xi^64,65^, and *Pola1*, a gene important for DNA replication during cell division^72^, were upregulated in GL7+ B cells from pristane-treated Xist cKO mice. Interestingly, *Xist* loss had minimal impact on X-linked expression in CD23+ B cells and plasma cells. In the case of CD23+ B cells, this is likely due to the heterogeneity of the CD23+ population, which may mask changes in expression from the Xi within select CD23+ B cell subsets. However, the absence of transcriptional changes in plasma cells, despite plasma cell accumulation in Xist cKO mice with spontaneous or induced disease, suggest that *Xist* loss influences differentiation into the plasma cell lineage but has less of an effect on plasma cell transcriptional status itself.

In addition to changes in X-linked gene expression, we identified autosomal transcriptional changes in Xist cKO B cells with SLE phenotypes. In CD11c+ ABCs from Xist cKO High mice, upregulated autosomal genes were enriched for genes involved in lymphocyte activation, including *Bcl3* (important for antigen-specific autoantibody formation)^73^, *Bmpr1a* (upregulated in some memory B cell subsets)^74^, *Cxcr4* (plays a role in B cell homeostasis and trafficking)^75^, and *Fas* (upregulated in ABCs from patients with SLE)^41^. In CD11c+ ABCs from pristane-treated Xist cKO mice, upregulated autosomal genes were enriched for genes involved in immune processes, including *Pik3r6* (mediates signaling pathways that regulate proliferation, class-switch recombination, and plasma cell differentiation)^76^, *Sh3kbp1* (poises B cells to respond to BCR stimulation)^77^, and *Zbtb20* (involved in plasma cell differentiation via Irf4)^78^. Unlike ABCs, upregulated autosomal genes in GL7+ B cells were enriched for genes involved in DNA replication and mitotic cell processes, including *Tipin*^79^, *Ska2*^80^, and *Mastl*^81^. These autosomal transcriptional changes, along with upregulation of X-linked linked genes including *Tasl*, likely underly B cell subset changes in Xist cKO mice and contribute to SLE phenotypes. GL7+ B cells, which are expanded in Xist cKO mice with spontaneous disease and expand as early as 4 months post pristane-injection, are characterized by transcriptional changes involved in cell proliferation. Meanwhile, CD11c+ ABCs, which are expanded in Xist cKO mice with spontaneous disease but do not expand until 7-9 months post-pristane injection, have transcriptional changes related to B cell differentiation and activation. These findings provide temporal clues as to how SLE-like disease develops in Xist cKO mice, and suggest that *Xist* deletion in the B cell compartment first drives germinal center reactions via proliferation and replication pathways, and later favors extrafollicular reactions that result in the accumulation of short-lived plasma cells and ABCs^82^.

Recent work has demonstrated that Xist RNA itself can contribute to the development of female-biased autoimmune disease. In these studies, the conserved repetitive elements at the 5’ end of Xist RNA were found to be strong ligands for TLR7 signaling, and XIST RNA oligos encompassing these motifs could stimulate type I interferon production *in vitro* by primary human plasmacytoid dendritic cells. SLE patients have higher levels of XIST RNA in PBMCs and B cells, and *XIST* expression in PBMCs in positively correlated with SLEDAI score in SLE patients^83^. Furthermore, XIST RNA binds many RNA binding proteins, and recent evidence suggests that the association of these XIST-binding proteins with the XIST RNA scaffold may provide a unique, female-biased source of autoantigens in the context of female-biased autoimmune diseases^84^. These findings suggest an intriguing, complementary role for *Xist* as a driver of autoimmune responses independent of its role in X-linked dosage compensation in some cell types. Importantly, our work demonstrates that disrupting the role of *Xist* for dosage compensation in B cells is sufficient to cause spontaneous SLE-like disease and also exacerbate pristane-induced disease. These studies, taken together with our findings, support critical complementary roles for the Xi, in both the presence and absence of *Xist*, that collectively and perhaps synergistically contribute to the development of female-biased autoimmunity. Thus, our findings suggest that disruption of proper maintenance of XCI in the B cell compartment, a potential mechanism at play in female patients with SLE who have disrupted patterns of XCI in B cells, likely contributes to female-biased disease phenotypes via dosage imbalances of pathogenic X-linked genes.

## ACKNOWLEDGEMENTS

We would like to thank the Penn and CHOP Flow Cytometry Cores for FACS and flow cytometry support; S. Prouty and the Penn Skin Biology and Diseases Resource-based Center (SBDRC) for tissue processing; the Penn Vet Comparative Pathology Core for pathologic assessment; D. Cutillo and D. Beiting and PennVet CHMI for library preparation and RNA sequencing; I. Raman and the Microarray Core Facility, University of Texas Southwestern Medical Center at Dallas for Autoantigen Microarray preparation; C. Syrett for initial characterization of Xist +/cKO mice; L. King for feedback on the manucript, and all members of the Anguera lab for insightful discussions. This research was supported by NIH R01 AI134834 and R01 AI168047 (to M.C.A.), Penn MSTP T32 Training Grant GM07170 (to C.D.L.), Rheumatology Research Foundation Future Physician Scientist Award (to C.D.L.), the Lupus Foundation of America Goldie Simon and Gina M. Finzi Awards (to C.D.L.), and the National Institutes of Health NIAID F30 AI174437 (to C.D.L.), Rheumatology Research Foundation Scientist Development Award (to N.J.), Scleroderma Research Foundation Postdoctoral Fellowship Award (to N.J.), Penn I3H and Colton Center for Autoimmunity Pilot Grant (to N.J.), and NIH R21-AR081588-01 (to M.C.A).

## AUTHOR CONTRIBUTIONS

Conceptualization: C.D.L., M.C.A., Methodology: C.D.L., M.C.A., Analysis: C.D.L., M.C.A., Investigation: C.D.L., M.C.A., Manuscript Writing: C.D.L., M.C.A., Manuscript Review & Editing: C.D.L., M.C.A., N.J., M.P.C., Funding Acquisition: C.D.L., M.C.A., N.J.

## DECLARATION OF INTERESTS

The authors declare no competing interests.

## SUPPLEMENTAL INFORMATION

Document S1. Figures S1-S6

Document S2. Tables S1-S9

## METHODS

### Mice

Xist^2lox^/Xist^2lox^ mice (129Sv/Jae strain) were a gift from Rudolf Jaenische and obtained from JeannieT.Lee^85^. Xist^2lox^/Xist^2lox^ mice were backcrossed onto the C57BL/6 or BALB/c background for at least 10 generations. Mb1-cre mice^86^ (B6.C(Cg)-Cd79^atm1(cre)Reth^/EhobJ; Jackson Laboratory Strain 020505) were purchased from Jackson Laboratory. Mb1^WT/cre^ mice were backcrossed onto the BALB/C background for at least 10 generations. To generate B-cell specific heterozygous *Xist* deletion (Xist +/cKO) mice, female Xist^2lox^/Xist^2lox^ mice were crossed to male mb1-cre^WT/cre^ mice. To generate B-cell specific homozygous *Xist* deletion (Xist cKO/cKO) mice, female mb1-cre^WT/cre^ Xist^2lox^/Xist^2lox^ mice were crossed to male mb1-cre^WT/WT^ Xist^2lox^/Y mice; reciprocal crosses were also performed using male mb1-cre^WT/cre^ Xist^2lox^/Y and female mb1-cre^WT/WT^ Xist^2lox^/Xist^1lox^ mice (Xist +/cKO). Mb1-cre was maintained heterozygous in every mating. PCR genotyping primers for mb1^WT^, mb1^cre^, Xist^WT^, and Xist^2lox^ alleles: (Mb1 Common: ACTGACGCAGGAGGATTGG, Mb1 Wildtype Forward (WT F): CTCTTTACCTTTCCAAGCACTGA, Mb1 Mutant Forward (mut F): CATTTTCGAGGGAGCTTCA; mb1^cre^ amplicon is 230bp, mb1^WT^ amplicon is 197bp) and Xist (Xprom F: TCTTCCCACTCTACCCTTGC, Xprom R: GACCAGAGTTCAAGTCCCCA, Xint3 R: CACTGGCAAGGTGAATAGCA; Xist^2lox^ amplicon is 500bp, Xist^1lox^ amplicon is 350bp, and Xist^WT^ amplicon is 300bp) primer sets. Animal experiments were approved by the University of Pennsylvania Animal Care and Use Committee (IACUC).

### Flow cytometry and cell sorting

Spleens and/or bone marrow were isolated from mice following euthanasia and flow cytometry was performed as previously described^23^. Splenocytes were isolated via mechanical dissociation of the spleen. Cell suspensions of the bone marrow were isolated via mechanical dissociation of the tibia and femur. Terminal serum was obtained from whole blood obtained by cardiac puncture. Red blood cells were removed by red blood cell lysis with ACK Lysis Buffer (Quality Biological 118-156-721). Cells were stained with LIVE/DEAD Fixable Aqua Dead Cell Stain Kit (Life Technologies L34965) and subsequently blocked with Purified Rat Anti-Mouse CD16/CD32 (BD Biosciences 553142). Cells were then stained in 0.5% BSA in PBS using the following fluorochrome-conjugated antibodies (clone, catalog #) purchased from BioLegend B220-APC (RA3-6B2, 103212), B220-BV650 (RA3-6B2, 103241), CD19-BV785 (6D5, 115543), CD23-BV421 (B3B4, 101621), CD21/35-APC-Cy7 (7E9, 123418), CD43-PE (S11, 143205), IgD-BV711 (11-26c.2a, 405731), GL7-PE-Cy7 (GL7, 144620), CD138-BV605 (281-2, 142516), CD11c-AF488 (N418, 117311), F4/80-PE-Cy5 (BM8, 123111), Gr-1-PE-Cy5 (RB6-8C5, 108410), CD4-PE-Cy5 (H129,19, 130312), CD8a-PE-Cy5 (53-6.7, 100709), or eBioscience CD93 (AA4.1, 25-5892-82), or BD Horizon IgM-PE-CF594 (R6-60.2, 562565). Flow cytometry was performed on an LSR Fortessa and sorting was performed on a FACSAria Fusion and data were analyzed using FlowJo software.

### Xist RNA fluorescent in situ hybridization (FISH) and Immunofluorescence (IF)

Naive splenic B cells were isolated from mice as previously described^26^. Briefly, spleens were mechanically homogenized and splenocytes were incubated with Biotin anti-mouse CD23 (Biolegend 101604 clone B3B4). Splenocytes were then incubated with Streptavidin MicroBeads (Miltenyi 130-048-101) and passed through a magnetic LS Column (Miltenyi 130-042-401). B cell activation was induced using 24-hour incubation with the Tlr9 agonist CpG (Invivogen tlrl-1826-1). *In vitro* activated lymphocytes were collected and prepared for FISH and IF as previously described^22^. Xist RNA FISH was performed using Cy3-labeled oligonucleotide probes for Xist RNA^22,23^, and nuclei were counterstained using Vectashield with DAPI (Vector Lab H-1200). Sequential immunofluorescence for heterochromatic histone marks was performed using primary antibodies for H2AUb119K (Cell Signaling 8240S) and secondary antibodies AF488-conjugated goat anti-rabbit IgG (Abcam ab150077-500UG). Samples were imaged using Nikon Eclipse Microscope and patterns of Xist RNA and histone focal enrichment were quantified manually as previously described^19^.

### Enzyme-linked immunosorbent assay (ELISA)

Blood was collected from mice using cheek bleeding procedure, and serum was isolated by centrifugation of blood that had been incubated at 4°C for 1 hour. ELISA was performed to measure serum levels of antibodies against double-stranded dsDNA, Smith-RNP, or IgG. Immulon 2 Flat Bottom High Binding 96 Well Plates (Fisher 1424561) were coated with calf thymus dsDNA (Alpha Diagnostics DNAD25-N-1), Sm-RNP (Arotec Diagnostics ATR01), or Total Ig (Southern Biotech 1010-01) in bicarbonate buffer (Sigma C3041-100). Plates were incubated overnight and then washed with ELISA Wash Buffer (1x PBS + 0.1% Tween) and blocked overnight with 1% BSA in PBS. Serum was diluted 1:200 - 1:1000 in 1% BSA in PBS. A standard curve was generated using mixed serum from mice with spontaneous SLE-like disease (NZB/W F1^25^, MRL/lpr ^26^, and NZM2328 mice^26^) diluted 1:500 in 1% BSA in PBS. Serum from mb1^cre/cre^ homozygous mice (which lack B cell development and do not generate antibodies) was used as a negative control. Serum samples were incubated overnight. HRP-conjugated goat anti-mouse IgG (Southern Biotech 1038-05) secondary antibody was added and incubated for one hour at room temperature, and SureBlue TMB 1-Component Microwell Peroxidase Substrate (LGC Clinical Diagnostics 5120-0075) was added for 10-25 minutes. TMB Stop Solution (LGC Clinical Diagnostics 5150-0020) was added to stop the reaction and plates were read using a plate reader at 450nm.

### Autoantibody profiling

Serum samples were submitted to the Genomics and Microarray Core Facility at the University of Texas Southwestern. Serum was analyzed for 120 autoantibodies and a mouse Ig control at 4 dilutions according to established methods. For each sample, a signal to noise ratio (SNR) was calculated to reveal high background, and autoantibodies where more than 50% samples per group had a SNR <3 were removed from analysis (6 autoantibodies). A “Net Signal Intensity” (NSI) value was reported by the core facility for each autoantibody which was then normalized to total IgG as measured by ELISA (“NSI_IgG”). An antibody score was calculated using the equation Antibody score = log2([NSI_IgG*SNR]+1). DESeq2 was used to identify significantly differentially enriched autoantibodies (“DEAs”) between groups. DEAs were filtered based on a Benjamini-Hochberg adjusted *p* value < 0.05.

### Proteinuria

Urine was collected from live mice and protein levels in the urine were measured by Pierce-Coomassie (Thermo Fisher Scientific 23200) according to manufacturer’s instructions.

### Enzyme-linked immunosorbent spot (ELISPOT)

ELISPOT assays were used to quantify numbers of anti-dsDNA producing splenocytes and bone marrow cells. 96-well MultiScreen HTS Plates (Millipore-Sigma MSIPN4550) were coated with 0.01% poly-L-lysine (Millipore Sigma A-005-C) for one hour and then coated overnight with calf thymus dsDNA (Alpha Diagnostics DNAD25-N-1) in bicarbonate buffer (Sigma C3041-100). Total splenocytes or bone marrow cells were plated at three dilutions in B cell media (RPMI containing 10% FBS, 1% nonessential amino acids, 1% 1M HEPES, 1% 200mM L-glutamine, 1% OPI medium supplement Hybri-Max, 1% Pen/Strep, and 0.1% β-mercaptoethanol) and plates were incubated overnight. Alkaline phosphatase conjugated goat anti-mouse IgG secondary antibody (Southern Biotech 1030-04) was added and incubated for two hours at room temperature, and BCIP/NBT Substrate (Thermo Pierce 34042) was added for 15 minutes. Plates were washed with water to stop the reaction and plates were imaged and analyzed using a CTL Immunospot Cell Imager and Analyzer.

### RT-qPCR

RT-qPCR was performed as previously described^26^. RNA was isolated from CD23+ splenic B cells in TRIzol and quantified using a NanoDrop™ spectrophotometer. cDNA was generated with qScript DNA SuperMix (QuantaBio), and qPCR was performed using PerfeCTa SYBR Green SuperMix (QuantaBio). Fold change expression was quantified by (2^−ΔΔCT^) using the CT value of *Xist* and the CT value of the housekeeping gene *Rpl13a*. The following primer sets were used: *Xist* (F: CAGAGTAGCGAGGACTTGAAGAG; R: GCTGGTTCGTCTATCTTGTGGG); *Rpl13a* (F: AGCCTACCAGAAAGTTTGCTTAC; R: GCTTCTTCTTCCGATAGTGCATC).

### RNA Sequencing

RNA was isolated from fluorescence-activated cell sorting (FACS) isolated populations of splenic B cells using the Qiagen RNeasy Micro Kit (Qiagen 74004). Library preparation of RNA samples was performed using the SMARTer Stranded Total RNA-Seq Kit v3 - Pico Input Mammalian (Takara 634485). Two independent pooled libraries (samples from mice with spontaneous disease and samples from mice with pristane-induced disease) were sequenced on 100 cycle flow cells using an Illumina NextSeq 2000 sequencer. Reads were aligned to a BALB/cJ reference genome (Ensembl BALB_cJ_v1) using STAR (v2.7.1a)^87^. Read counts were calculated using featureCounts^88^. RNAseq data was analyzed using RStudio (v1.2.5042). edgeR (v3.32.1) was used to filter data and data was normalized using quantile normalization^89^. Differentially expressed genes were identified using DESeq2 (v1.30.1). Pathway analysis was performed using Metascape^90^. Venn diagrams were generated using DeepVenn.

### Kidney Histopathology

Kidneys were isolated from euthanized mice and formalin fixed for 24 hours (Sigma-Aldrich HT501128-4L). Formalin-fixed kidneys were processed by the Penn Skin Biology and Diseases Resource-based Center: kidneys were embedded in paraffin, and organ sections were stained with hematoxylin and eosin (H&E) or periodic acid-Schiff (PAS). Stained sections were scored for features of kidney disease by a blinded veterinary renal pathologist who measured features of kidney disease (glomerular hypercellularity, glomerular mesangial thickening, glomerular cystic dilation, glomerular basement membrane thickening, glomerular hypertrophy, glomerular mesangiolysis, glomerulosclerosis, tubular basophilia/regeneration, tubular degeneration, tubular dilation, and tubular casts). Each feature was scored from a range of 0-4: 0 = “no significant findings”, 1 = “minimal findings”, 2 = “mild findings”, 3 = “moderate findings”, and 4 = “severe findings”.

### Pristane Treatment

2,6,10,14-Tetramethylpentadecane (“pristane”) (Sigma Aldrich P2870) was administered to mice for the induction of SLE-like disease as previously described^91^. Briefly, 500uL of pristane was injected intraperitoneally into 3-5 month old female Xist cKO and WT mice on the BALB/c background. Seven independent cohorts of mice were designed with designated experimental endpoint set at onset (4 months, 7 months, or 9 months). Mice were monitored over the course of 0-9 months for serum dsDNA autoantibodies by ELISA and proteinuria. Mice were euthanized at the pre-set experimental endpoint.

### Statistical Analysis

Serum autoantibodies, kidney pathology scores, and B cell subsets frequencies and total numbers were compared between multiple groups using an ordinary one-way ANOVA with Tukey’s multiple comparisons test or compared between two groups using Mann-Whitney U tests for non-normally-distributed data. Statistical significance was defined as *p* < 0.05. Differentially enriched autoantibodies were defined using DESeq2 to identify autoantibodies with a Benjamini-Hochberg-adjusted *p*-value (FDR) < 0.05. Changes in serum autoantibodies or proteinuria over pristane-induced disease between groups were compared using a mixed effects model with the Geisser-Greenhouse correction (not assuming equal variability of differences). For RNAseq analysis, genes exhibiting a fold-change ≥0.5 with a Benjamini-Hochberg-adjusted *p*-value (FDR) < 0.05 were considered differentially expressed. The total number of animals used for each experiment is indicated in the corresponding figure legend. Data are pooled from multiple independent experiments.

### Data Availabilty

All sequencing data generated in this study has been deposited to the NCBI GEO database. Access data using the following accession numbers: GSE266417, GSE266231.

## SUPPLEMENTAL FIGURE AND TABLE LEGENDS

**Figure S1. Xist cKO mice lack *Xist* expression in the B cell compartment and maintain intact B cell development**

**A.** Representative images from sequential Xist RNA fluorescent in situ hybridization (FISH) and H2AK119Ub immunofluorescence (IF) from splenic CD23+ B cells stimulated with CpG for 24 hours from WT, Xist +/cKO, and Xist cKO/cKO mice. **B.** Quantification of the frequency of nuclei with a localized Xist RNA focus based on Xist RNA FISH (n=3 biological samples quantified per genotype). Data are depicted as the mean percentage of nuclei with an Xist RNA focus +/- SD. **C.** Quantification of the frequency of nuclei with an H2AK119Ub focus (n=3 biological samples quantified per genotype). Data are depicted as the mean percentage of nuclei with an H2AK119Ub focus +/- SD. **D.** Xist RNA expression as measured by qPCR. Fold-change per sample (n=3 Xist +/cKO; n=8 Xist cKO/cKO) relative to average expression of the WT samples (n=8). **E.** Survival of n=17 female WT mice, n=22 female Xist +/cKO mice, and n=11 female Xist cKO/cKO mice. “ns” = not statistically significant by log-rank (Mantel-Cox) test. **F.** Quantification of splenic B cells (B220+ CD19+). **G.** Quantification of bone marrow B cells (B220+). * = *p* <0.05, ** = *p* < 0.01, *** = *p* < 0.001, **** = *p* < 0.0001 via one-way ANOVA with Tukey’s multiple comparisons test unless otherwise specified.

**Figure S2. Some Xist cKO/cKO and Xist +/cKO mice develop serologic features of SLE**

**A.** Serum anti-dsDNA autoantibodies in n=109 WT, n=45 Xist +/cKO, and n=66 Xist cKO/cKO female BALB/c mice 6 weeks - 18 months old and n=8 NZB/W mice. **B.** Timeline of female BALB/c development of elevated anti-dsDNA autoantibodies. Each dot represents a single mouse at a given age. Any mouse that developed anti-dsDNA at or above levels of NZB/W mice are shown as an uptick in the line at the age at which that mouse developed elevated anti-dsDNA. The y-axis depicts the frequency of mice with high anti-dsDNA autoantibodies relative to the total number of available mice at that age. **C.** Number of anti-dsDNA producing cells per 1,000,000 bone marrow cells as measured by ELISPOT. **D.** Total serum IgG as measured by ELISA. **E.** Representative anti-dsDNA ELISPOT images from WT, Xist cKO Low, and Xist cKO High splenocytes. **F.** Representative anti-dsDNA ELISPOT images from WT, Xist cKO Low, and Xist cKO High bone marrow. **G.** Autoantibodies significantly lower in Xist cKO High mice compared to WT mice or Xist cKO mice. **H.** Autoantibodies significantly higher in Xist cKO High mice compared to WT mice or Xist cKO mice. **I-O.** All cells were pre-gated on live, single CD4-CD8-Gr-1-F4/80- cells. **I.** Splenic B cells (B220+ CD19+). **J.** Bone marrow B cells (B220+). **K.** Splenic GL7+ activated B cells (B220+CD19+IgM-IgD-GL7+)**. L.** Splenic class-switched B cells (B220+CD19+IgM-IgD-). **M.** Splenic short-lived plasma cells (“SLPCs”; IgD-CD138+B220+) **N**. Splenic age-associated B cells (“ABCs”; B220+CD19+CD11c+). **O.** Splenic long-lived plasma cells (“LLPCs”; IgD-CD138+B220-). **P.** Bone marrow short-lived plasma cells (“SLPCs”; IgD-CD138+B220+) and long-lived plasma cells (“LLPCs”; IgD-CD138+B220-). Data in bar graphs are depicted as mean ± SD. * *p* < 0.05, ** *p* < 0.01, *** *p* < 0.001, **** *p* < 0.0001 via ordinary one-way ANOVA with Tukey’s multiple comparisons test unless otherwise specified.

**Figure S3. B cells from Xist cKO High mice demonstrate genome-wide transcriptional changes**

**A.** Principal component analysis of RNA sequencing data from splenic CD23+ B cells from Xist cKO High, Xist cKO Low, and WT mice. **B.** *Xist* RNA expression in splenic CD23+ B cells from WT, Xist cKO Low, and Xist cKO High mice, reported as counts per million. **C.** Heatmap of differentially expressed autosomal genes between Xist cKO High and WT splenic CD23+ B cells. **D.** Frequency of GL7+ activated B cells as a frequency of CD23+ splenotyes. ** *p* < 0.01 via ordinary one-way ANOVA with Tukey’s multiple comparisons test. **E.** Principal component analysis of RNAseq data from splenic CD11c+ ABCs from Xist cKO High mice and WT controls. **F.** *Xist* RNA expression in splenic CD11c+ ABCs from WT and Xist cKO High mice reported as counts per million. **G.** *Tlr7* expression in splenic CD11c+ ABCs from WT and Xist cKO High mice reported as counts per million. **H.** Heatmap of differentially expressed autosomal genes between Xist cKO High and WT CD11c+ ABCs. **I.** Principal component analysis of RNAseq data from SLPCs from Xist cKO High mice and WT controls.

**Figure S4. Pristane-treated Xist cKO mice share some disease features with pristane-treated WT mice**

**A.** Mean proteinuria levels ± SEM in pristane-treated Xist cKO mice (n=12), pristane-treated WT mice (n=16), and PBS-treated WT mice (n=3) over 7 months. Not significant by mixed effects model with the Geisser-Greenhouse correction (not assuming equal variability of differences). **B-C.** All cells were pre-gated on live, single CD4-CD8-Gr-1-F4/80-cells. Quantification of splenic (**B**) B cells (B220+) or (**C**) long-lived plasma cells (“LLPCs”; IgD-CD138+B220-) after 4, 7, or 9 months post-pristane. Data are presented as mean ± SD. * = *p* < 0.05 via Mann Whitney test comparing pristane-treated Xist cKO to pristane-treated WT.

**Figure S5. Pristane-treated Xist +/cKO mice develop an intermediate SLE phenotype**

**A.** Pristane-induced SLE experimental design. Female WT and Xist +/cKO mice were injected intraperitoneally with pristane and mice were monitored for up to 9 months for development of elevated autoantibodies in the blood and proteinuria. At experimental endpoint, organs were harvested for flow cytometry or pathologic assessment. N=4 WT mice and n=2 Xist +/cKO mice died spontaneously during disease development, and are not included in Endpoint cohorts. **B.** Mean serum anti-dsDNA autoantibody levels ± SEM in pristane-treated Xist +/cKO mice (n=20), pristane-treated WT mice (n=41), and PBS-treated WT mice (n=10) over 7 months. Not significant by mixed effects model with the Geisser-Greenhouse correction (not assuming equal variability of differences). **C.** Mean serum anti-dsDNA autoantibody levels ± SD in pristane-treated Xist +/cKO mice (n=6), pristane-treated WT mice (n=18), and PBS-treated WT mice (n=3) at 9 months post-pristane injection. **D.** Mean proteinuria levels ± SEM in pristane-treated Xist +/cKO mice (n=11), pristane-treated WT mice (n=16), and PBS-treated WT mice (n=3) over 9 months. Not significant by mixed effects model with the Geisser-Greenhouse correction (not assuming equal variability of differences). **E.** Mean total glomerular pathology score ± SD. **F.** Mean total tubular pathology score ± SD. **G.** Mean interstitial inflammation ± SD. **H-M.** All cells were pre-gated on live, single, CD4-CD8-Gr1-F4/80-cells. Quantification of splenic (**H**) total B cells (B220+), (**I**) class-switched B cells (B220+CD19+IgM-IgD-), (**J**) GL7+ activated B cells (B220+CD19+IgM-IgD-CD138-GL7+), (**K**) short-lived plasma cells (SLPCs; IgD-CD138+B220+), (**L**) long-lived plasma cells (LLPCs; IgD-CD138+B220-), and (**M**) age-associated B cells (ABCs; B220+CD19+CD11c+) after 4, 7, or 9 months of pristane. Data are presented as mean ± SD. * *p* < 0.05, ** *p* < 0.01 via Mann-Whitney U test comparing pristane-treated Xist cKO to pristane-treated WT unless otherwise specified.

**Figure S6. Activated B cells from pristane treated Xist cKO mice demonstrate marked autosomal transcriptional changes**

**A.** Principal component analysis (PCA) of RNAseq from WT and Xist cKO pristane ABCs. **B.** Heatmap of differentially expressed autosomal genes in CD11c+ ABCs from pristane-treated mice. **C.** PCA of RNAseq from WT and Xist cKO pristane GL7+ activated B cells. **D.** *Xist* RNA expression as measured by counts per million in GL7+ B cells from pristane treated WT and Xist cKO mice. **E.** Heatmap of differentially expressed autosomal genes in GL7+ activated B cells from pristane-treated mice. **F.** PCA of RNAseq from WT and Xist cKO pristane SLPCs.

**Table S1. Autoantibody scores from autoantigen microarray**

**Table S2. Log2 counts per million of significantly differentially expressed genes (DEGs) in CD23+ B cells from Xist cKO High vs. WT mice**

**Table S3. Log2 counts per million of DEGs in CD23+ B cells from Xist cKO Low vs. WT mice**

**Table S4. Log2 counts per million of DEGs in CD11c+ ABCs from Xist cKO High vs. WT mice**

**Table S5. Log2 counts per million of DEGs in long-lived plasma cells from Xist cKO High vs. WT mice**

**Table S6. Log2 counts per million of DEGs in CD11c+ ABCs from pristane-treated WT and Xist cKO mice**

**Table S7. Log2 counts per million of DEGs in GL7+ B cells from pristane-treated WT and Xist cKO mice**

**Table S8. Log2 counts per million of DEGs in SLPCs from pristane-treated WT and Xist cKO mice**

**Table S9. Metascape pathway analysis from all RNA sequencing analyses**

## Notes

### Competing Interest Statement

The authors have declared no competing interest.

