## Supplemental Figures for "*Xist* Deletion in B Cells Results in Systemic Lupus Erythematosus Phenotypes": 20240513 Xist cKO Supplemental Figures Compressed.pdf

### Supplemental Figure 1

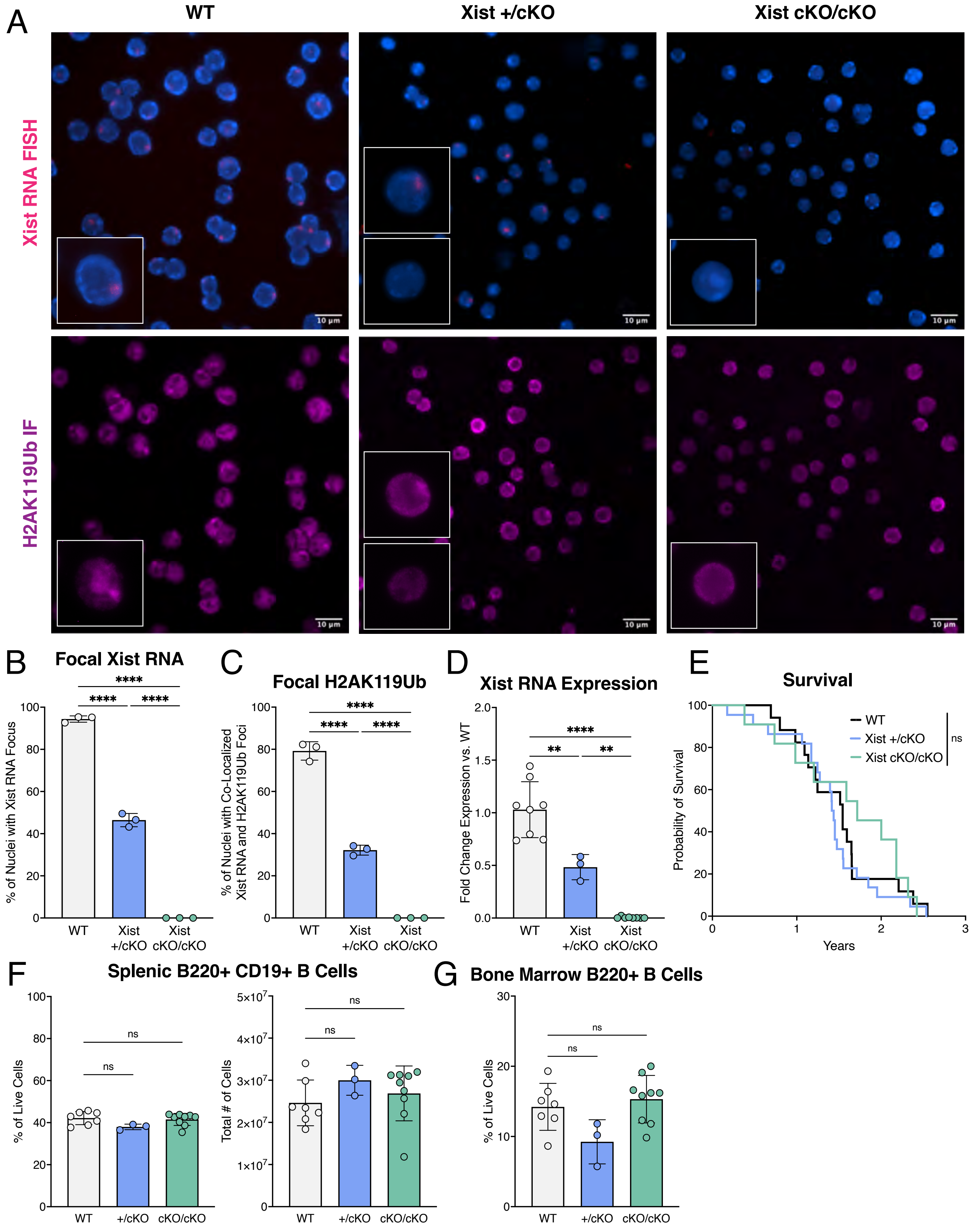

### Supplemental Figure 2

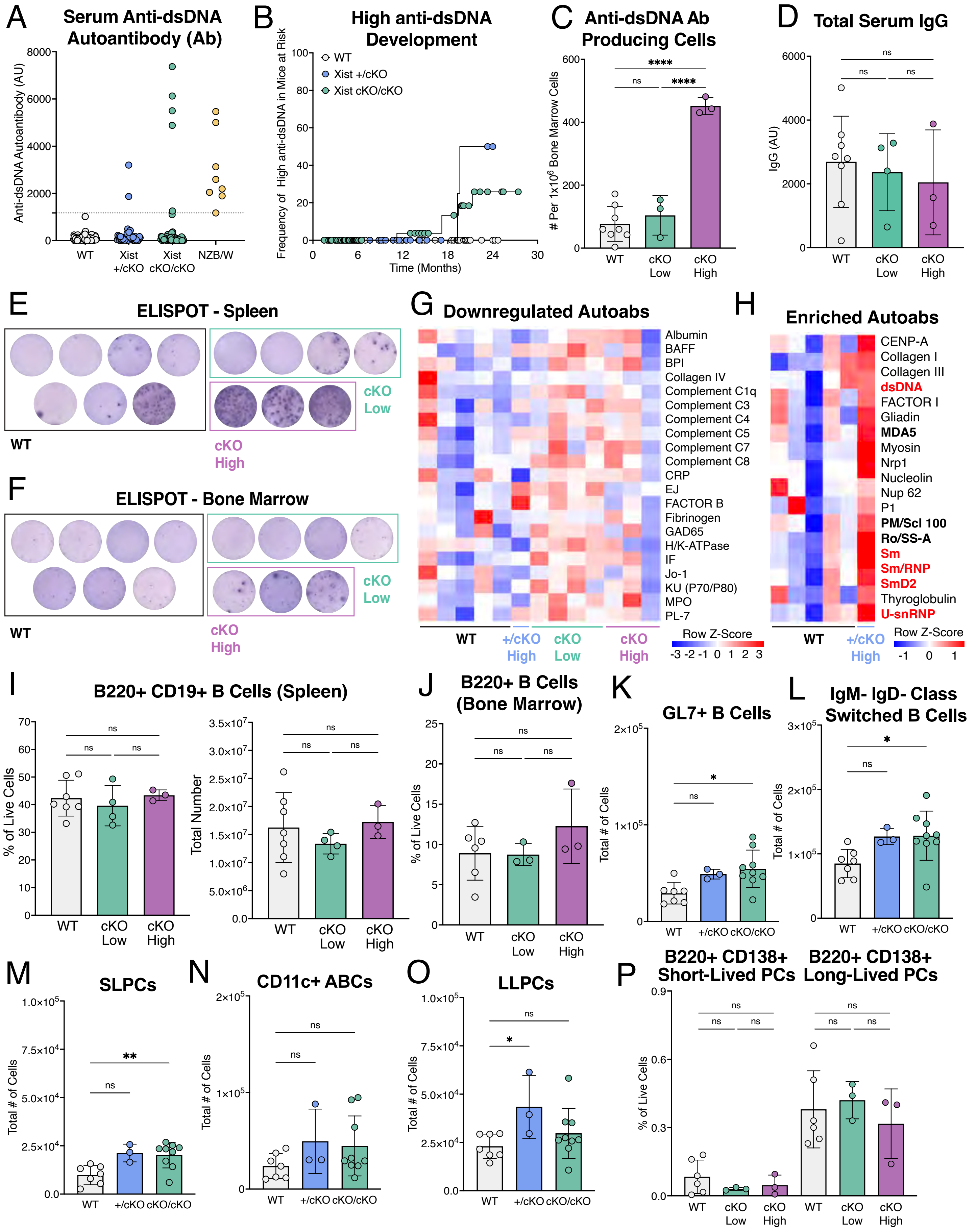

### Supplemental Figure 3

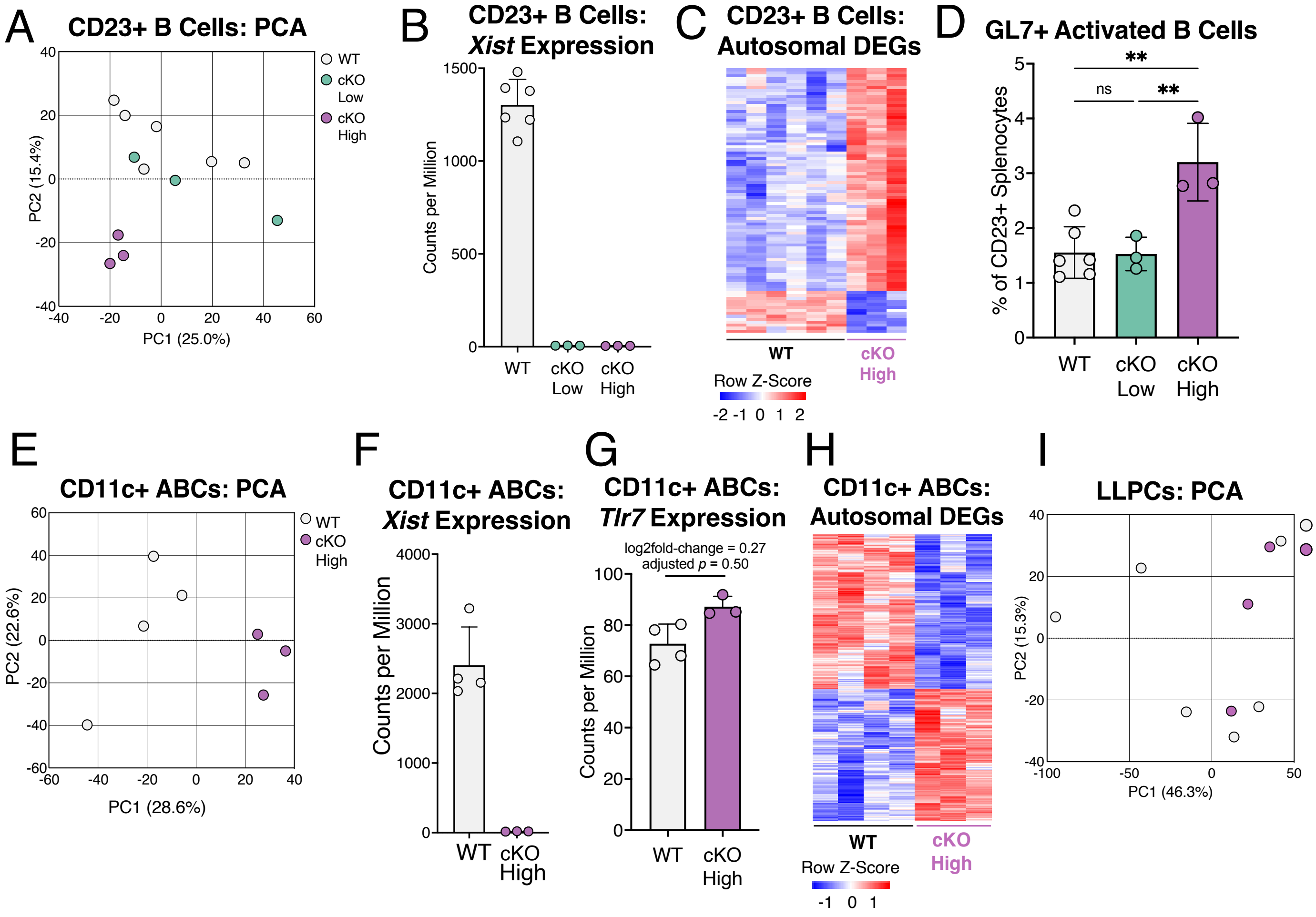

### Supplemental Figure 4

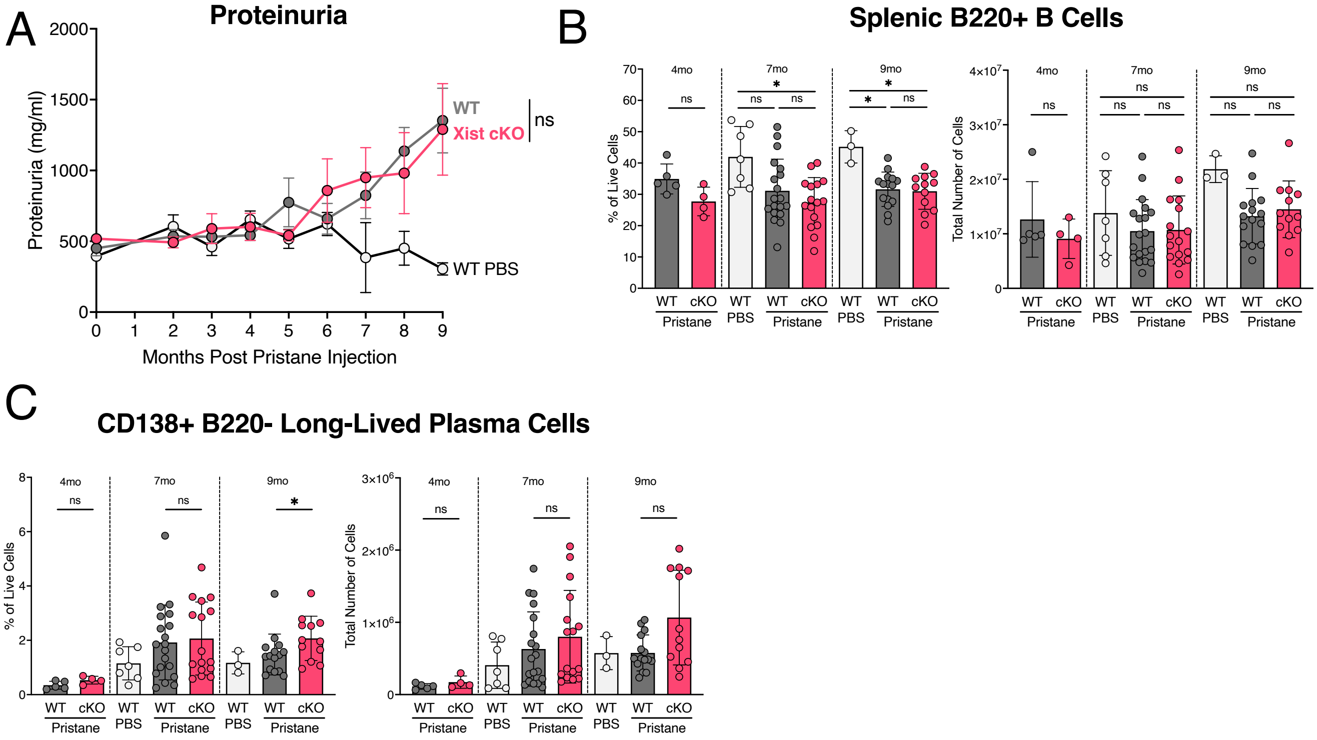

### Supplemental Figure 5

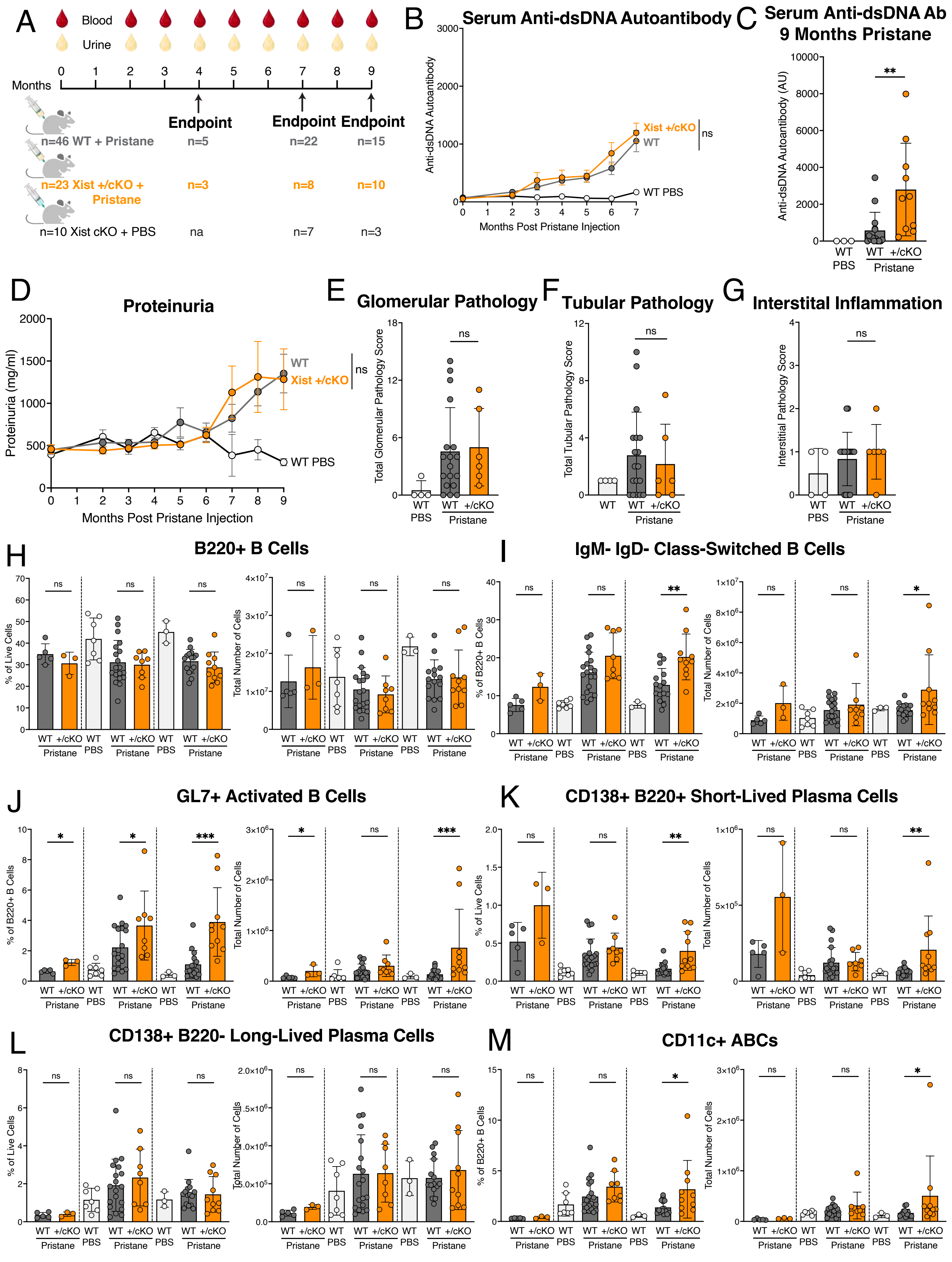

### Supplemental Figure 6

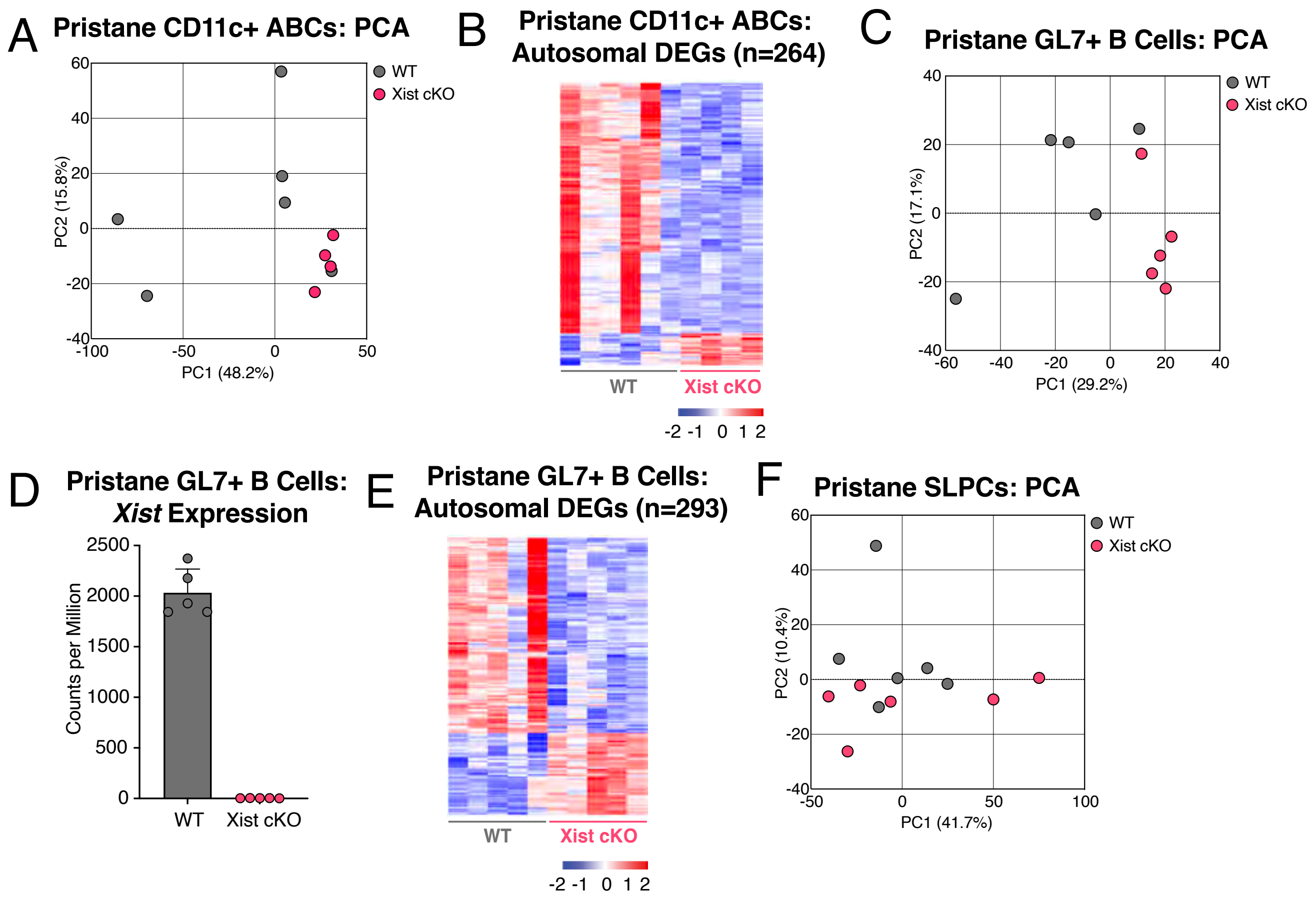
